# Coordinated wound responses in a regenerative animal-algal photosymbiotic metaorganism

**DOI:** 10.1101/2023.06.21.545945

**Authors:** Dania Nanes Sarfati, Yuan Xue, Eun Sun Song, Ashley Byrne, Daniel Le, Spyros Darmanis, Stephen R. Quake, Adrien Burlacot, James Sikes, Bo Wang

## Abstract

Animal regeneration requires coordinated responses of many cell types throughout the animal body. In animals carrying endosymbionts, cells from the other species may also participate in regeneration, but how cellular responses are integrated across species is yet to be unraveled. Here, we study the acoel *Convolutriloba longifissura*, which hosts symbiotic *Tetraselmis* green algae and can regenerate entire bodies from small tissue fragments. We show that animal injury leads to a decline in the photosynthetic efficiency of the symbiotic algae and concurrently induces upregulation of a cohort of photosynthesis-related genes. A deeply conserved animal transcription factor, *runt*, is induced after injury and required for the acoel regeneration. Knockdown of *runt* also dampens algal transcriptional responses to the host injury, particularly in photosynthesis related pathways, and results in further reduction of photosynthetic efficiency post-injury. Our results suggest that the *runt-*dependent animal regeneration program coordinates wound responses across the symbiotic partners and regulates photosynthetic carbon assimilation in this metaorganism.

## Introduction

Animal regeneration encompasses multiple cellular and molecular processes which require coordination across different cell types ^1–3^. For endosymbiotic animals, regeneration must be achieved in the presence of other species, adding an additional layer of complexity to the coordination of cellular responses ^4^. It has been increasingly recognized that symbiosis can profoundly modify morphogenesis and plasticity of the host tissues ^5,6^. In addition to examples in development, such as *Vibrio fischeri* symbionts guiding the formation of the squid’s light organ ^7,8^ and *Blochmannia* bacteria rewiring Hox genes during the embryonic development of Camponotini ants ^9^, the parallels between regeneration and developmental programs imply that symbiotic associations can also play important roles during regeneration. Indeed, studies have begun to elucidate the contributions of bacteria during regeneration of animals including sea cucumbers ^10^ and mice ^11^. However, less is known about how endosymbionts are regulated by the host regeneration and whether symbiont responses are integrated in the host regeneration program.

Progress in this respect has been bottlenecked by our limited capacity to measure the physiological states of symbionts. Here we overcome this challenge by studying animal-algae photosymbiosis. In this relationship, the animal host benefits from the photosynthetic capabilities of algal partners, which fix carbon into organic compounds by converting solar energy to chemical energy through the photosynthetic electron transport chain ^12^. Importantly, photosynthesis can serve as a gauge for algal physiology. In free living algae, photosynthesis is modulated by multiple abiotic factors, including light ^13^, nutrients ^14^, temperature ^15^, and the availability of inorganic carbon ^16^. In photosymbiosis, hosts can regulate the photosynthetic output of their endosymbiotic algae through modulating the concentrations of inorganic carbon and nitrogen ^17,18^, adjusting the pH of the symbionts’ microenvironment ^19,20^, and changing light intensity ^21^.

Specifically, we study the photosymbiotic acoel *Convolutriloba longifissura*, a marine worm that maintains an obligate symbiosis with *Tetraselmis* green algae ^22^. The algae reside between acoel cells, making this relationship an extracellular endosymbiosis ^23^. The acoels acquire their symbionts after hatching, and can transfer the algae to their progeny through asexual fission ^24,25^. *C. longifissura* fissions every few days, generating new individuals through regeneration of its entire body from tissue fragments ^24,26^. In contrast to embryonic development ^27^, regeneration proceeds in the presence of algal symbionts.

In this work, we assemble high-quality transcriptomes for both the acoel and the algae and develop a suite of tools to evaluate the host and endosymbionts’ responses during regeneration at the molecular and physiological levels. We find that, along with the expected acoel response, the algae exhibit an abrupt decrease in photosynthetic efficiency within the first few hours after host injury, accompanied by large scale transcriptional changes including the upregulation of pathways related to photosynthesis, carbon concentrating mechanisms, and chlorophyll biosynthesis. Notably, this contrasts the transcriptional changes induced by light stress in a similar timeframe, implicating that host injury triggers distinct algal responses that may be involved in specific requirements for acoel regeneration. In contrast, knockdown of a conserved injury-induced acoel transcription factor, *runt* ^28,29^, blocks acoel regeneration, reduces the algal transcriptional changes in photosynthesis-related genes, and further decreases the operating yield of photosystem II (PSII). As the function of *runt* during acoel regeneration appears to be conserved, our results suggest that the acoel’s early wound response contributes towards integrating the responses across species within this photosymbiotic metaorganism.

## Results

### *C. longifissura* anterior regeneration is epimorphic and does not require algal photosynthesis

*C. longifissura* belongs to a derived family of acoels (**Fig. 1a**), the Convolutidae, which is the only family of the Acoela taxon that has evolved endosymbiosis and robust asexual reproduction ^30^. The animal’s orange-red color is due to its red pigment cells and green chloroplasts within the algal symbionts (**Fig. 1b,c**). The chloroplasts, autofluorescent in the red spectrum, facilitate fluorescence imaging to identify algal distribution during homeostasis and regeneration (**Fig. 1d-f**). Algal cells are distributed throughout the acoel’s body, primarily accumulating beneath the body wall ^23^, which is a simple layer of epidermis lined with muscle and gland cells. A lower density of algal cells can also be found in the inner vacuolated parenchyma (**Fig. 1d**).

**Fig. 1:**
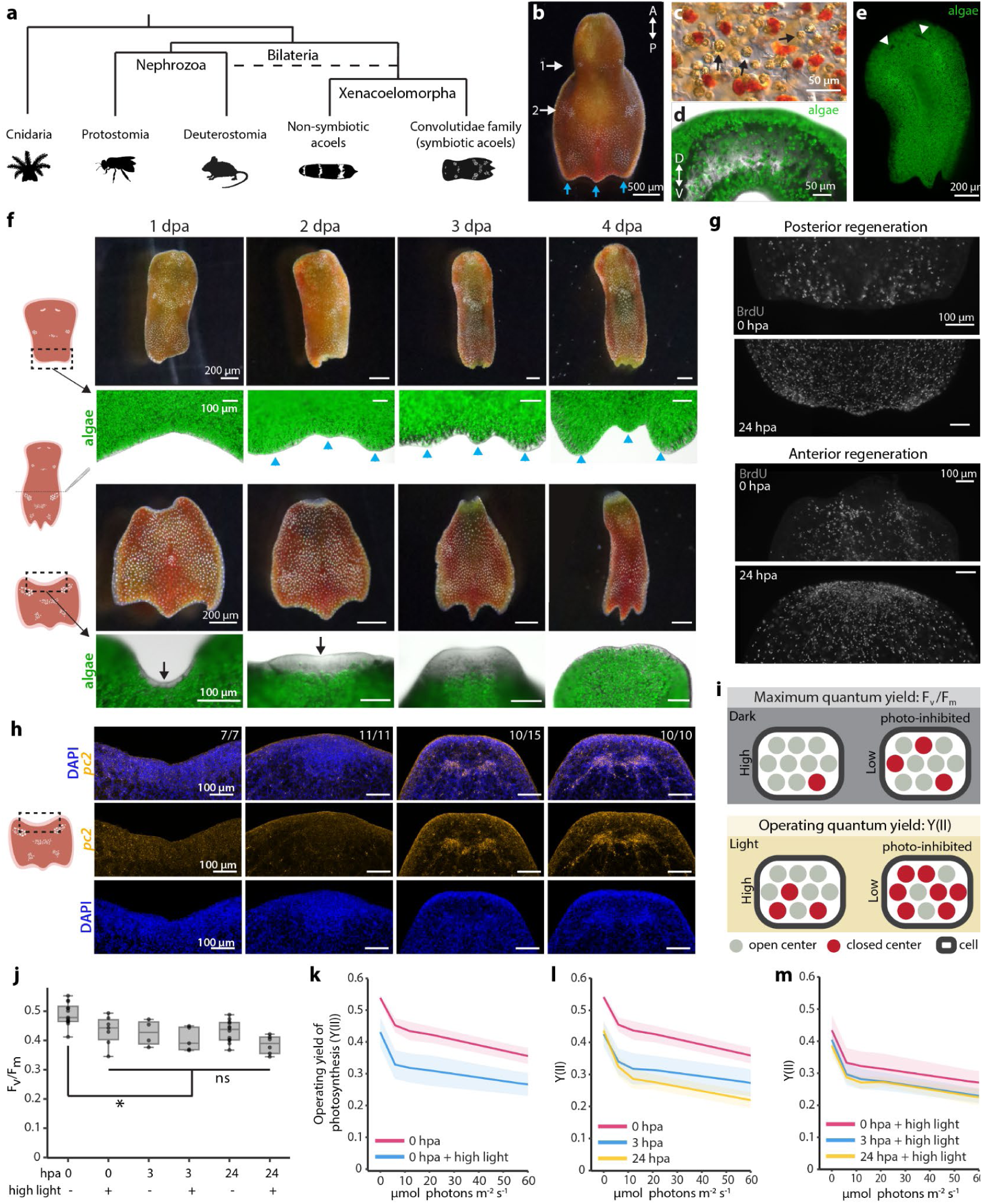
Acoel regeneration affects algal photosynthesis. **a** Schematic of a simplified phylogeny showing the position of symbiotic acoels, which belong exclusively to the Convolutidae family. The position of the Xenacoelomorpha is still debated ^83^ as a basal bilaterian or a basal deuterostome (dotted line). **b** Photograph of the acoel *C. longifissura*. Numbers mark the pairs of white concrement granules and blue arrows point to the three tail lobes. **c** Differential interference contrast image showing the acoel cells (transparent and red cells) intermingled with green-brown algal cells (a few examples are highlighted by arrows). **d** Transverse section at the second concrement granule pair showing the distribution algae (green) along the animal dorsal ventral axis (D-V), imaged through autofluorescence at 647 nm. **e** Algal cells are ubiquitously distributed across the host body, except the eye spots (arrowheads), which are devoid of algae. **f** Regeneration from posterior and anterior facing wounds. Top: posterior regeneration, with blue arrowheads indicating the regenerated tail lobes. Bottom: anterior regeneration with back arrows pointing at the blastema. Note the new tissue is devoid of algae until 4 dpa. **g** BrdU staining of posterior (top) and anterior (bottom) facing wound sites at 0 and 24 hpa. A buildup of BrdU^+^ cells is visible at the anterior wound at 24 hpa. **h** Fluorescence *in situ* hybridization of the neuronal marker *prohormone convertase 2* (*pc2*) and nuclei staining with DAPI at various time points post-amputation. The blastema formation is evident by the accumulation of cells at 2 dpa based on the DAPI staining. Neural ganglions are regenerated by 3 dpa. **i** Schematics showing the biological interpretation of F_v_/F_m_ and Y(II) measurements. At any given moment, a PSII can be available to receive electrons (open center) or unavailable (closed center). When a PSII is open and exposed to light, it can accept photons and use them for photochemistry. A closed center can be occupied or damaged. When a closed center receives light, the energy is transferred to an open center or dissipated as fluorescence or heat. Fluorescence is therefore used to measure changes in the ratio of open to close centers (an example trace is shown in **Supplementary Fig. 3**). F_v_/F_m_ is measured after dark acclimation, allowing all available, undamaged centers to open, and thus represents the maximum fraction of open centers. Y(II) is measured in the presence of actinic light, indicating the efficiency with which the electron transport chain makes use of electrons generated by PSII and thus open more centers to generate new electrons. If the photosynthetic electron transport chain gets saturated, PSII centers remain closed reducing Y(II). **j** Box plot showing F_v_/F_m_ of tails under different conditions: after certain hours post-amputation (hpa), with or without light stress. For 0 hpa samples, animals were dark acclimated for 20 min and then amputated under green light, to avoid stimulating photochemistry. For each treatment we conducted 4-13 biological replicates, each replicate consisted of 20 tails. Boxes represent upper and lower quartiles with the median marked by the middle line, the bars represent the upper and lower fences. The black dots show values of individual replicates. Treatments are significantly different compared to the ‘0 hpa’ data, whereas there is no significance among any other conditions (one-way ANOVA and a Tukey post hoc test). * p-adj ≦ 0.1, ns, no significance. **k-m** Y(II) measurements with increasing actinic light intensities of tail fragments with and without high light exposure for 24 hr before amputation (**k**), at different time points post amputation (**l**), and exposed to high light for 24 hr and then amputated (**m**). Each treatment had 3-7 biological replicates, each replicate consisted of 20 tails. The lines represent the mean values of different treatments. Shaded regions represent the area between one standard deviation above or below the mean.

The anterior of the *C. longifissura* body contains the neural ganglion and two eye spots, while the posterior pole is characterized by a three-lobed tail. Two pairs of white concrement granules, situated laterally, are present on the acoel’s dorsal surface: one posterior to the head and the second posterior to the gut-like syncytium ^31^ (**Fig. 1b**). The second pair of concrement granules coincides with the transverse fission plane ^24^, at which we bisect the animals to assess the wound responses and regeneration process.

The head fragments, undergoing posterior regeneration, do not form an obvious blastema. Newly formed tails regain their characteristic three-lobed morphology at 2 days post amputation (dpa, upper panels, **Fig. 1f**), similar to regeneration after fission ^24^. Fluorescence imaging reveals that algal cells persist in the posterior wound and regenerating tissues (upper panels, **Fig. 1f**). BrdU staining, which labels cells in S-phase and their progeny after division ^26^, indicates limited proliferation in the posterior wound region (**Fig. 1g**). These observations suggest that posterior regeneration follows a morphallactic process ^32^.

Conversely, during anterior regeneration from tail fragments, at 1 dpa BrdU^+^ cells accumulate towards the wound site (**Fig. 1g**) and form a clear blastema, distinguished by its transparency due to the absence of pigment and algal cells (lower panels, **Fig. 1f**). Accumulation of cells in the blastema is observable with DAPI staining at 2 dpa (**Fig. 1h**). The new tissue expands over the next couple of days, with head structures, including eye spots and the neural ganglion, restored by 4 dpa (**Fig. 1f, h, Supplementary Fig. 1**). In parallel, algae repopulate the new tissue between 3 and 4 dpa (lower panels, **Fig. 1f**). These features of anterior regeneration are consistent with epimorphosis ^32^. Given that anterior regeneration requires tissue growth and algal repopulation, we chose it as our focus for the rest of the study.

Despite the obligate photosymbiotic relationship, photosynthesis is not required for regeneration. Acoels can regenerate normally when kept in the dark throughout the regeneration process (**Supplementary Fig. 2**). Animals treated with 3-(3,4-dichlorophenyl)-1, 1-dimethylurea (DCMU), a chemical inhibitor of PSII, showed impaired PSII activity (**Supplementary Fig. 3**) but similar regeneration rates as controls. Longer DCMU treatment eliminated algal cells and led to animal death (**Supplementary Fig. 4**).

### Algal photosynthetic efficiency decreases upon acoel injury

Since algal photosynthesis is not essential for regeneration, we tested whether regeneration affected the symbiotic algae’s photosynthesis. For this, we designed a custom acoel chamber (**Supplementary Fig. 3**) mounted on a Pulse-Amplitude-Modulation (PAM) fluorometer. By measuring chlorophyll fluorescence changes we infer the ratio of PSII able to receive electrons for photochemistry (open centers) to the ones that are either electron-occupied or damaged (closed centers) ^33,34^ (**Fig. 1i**). The maximum quantum yield of PSII (F_v_/F_m_) measures the maximum fraction of open centers after a dark incubation as a proxy for photo-inhibition. The quantum yield of PSII (Y(II)) measures the fraction of open centers at any given time with background light, depicting the efficiency with which light energy is converted into chemical energy ^34,35^.

As a reference of algal photosynthesis in the host, we first established the effects of exposure to high light, which is known to lower F_v_/F_m_ of free-living algae ^36^. Excess light can overexcite the photosynthetic machinery, saturating the photosynthetic electron transport chain and producing reactive oxygen species (ROS), which may result in degradation of photosystems in a process called photo-damage ^37^. We quantified F_v_/F_m_ and Y(II) on tails (amputated and immediately measured), as whole animals moved too much to be measured reliably. We compared controls kept under constant light (“0 hpa”, hours post amputation) and acoels exposed to high light for 24 hr before measurement (labeled as “0 hpa + light stress”) (**Fig. 1j,k**). As expected, high light exposure caused a decrease in F_v_/F_m_ which was paralleled with a decrease in PSII efficiency for all actinic light intensities tested (**Fig. 1j,k**).

We then evaluated the effects of amputation on photosynthesis in a matched timeframe (0 and 24 hpa). We added a 3 hpa time point as animal wound responses are already pronounced at this early time ^28,29^, which may also induce changes in algal physiology. Surprisingly, both F_v_/F_m_ and Y(II) decreased concordantly at 3 hpa (**Fig. 1j**) but only Y(II) continued to decrease at 24 hpa (**Fig. 1l**). This may be caused by further interference with the electron transport chain’s photoconversion capacity at this later time point without affecting the maximum fraction of functional open centers. We conclude from this that upon amputation of the host, the algal photosynthetic capacity is decreased, likely reflecting PSII photo-inhibition.

Light-stressed animals at 3 or 24 hpa showed barely reduced F_v_/F_m_ beyond the effect of light stress alone (**Fig. 1j**), while Y(II) only had a minor decrease at 3 hpa without further reduction at 24 hpa (**Fig. 1m**). Both F_v_/F_m_ and Y(II) in amputated animals under light stress were comparable to values at 24 hpa with no light stress, suggesting that the effects of high light exposure and acoel amputation are non-additive.

### D*e novo* transcriptome assembly to assess molecular wound responses in acoel and algal cells

Given the similar degree of photosynthetic efficiency changes induced by injury and light, we hypothesized that host injury may induce molecular responses in the symbiotic alga, as observed in many microalgae subjected to light stress ^38^. To characterize gene expression changes, we needed reference transcriptomes that include genes expressed during homeostasis and after injury, which are lacking for both *C. longifissura* and *Tetraselmis* sp. Therefore, we assembled *de novo* transcriptomes using tissues collected throughout an early regeneration time course (**Fig. 2a, Supplementary Table 1**). To obtain full length mRNAs and enhance the quality of the assembly, we used Nanopore long-read sequencing to generate reads covering larger portions of transcripts with greater overlaps between fragments, which reduces ambiguity during assembly. However, Nanopore reads had frequent errors disrupting open reading frames (ORFs) of assembled transcripts. To address this, we polished the transcriptome using Pacbio ISO-Seq and Illumina short-read sequencing of the same cDNA (**Supplementary Fig. 5**). After removing duplicate and chimeric contigs, we obtained a final transcriptome consisting of 21,191 transcripts including both acoel and algal genes (**Supplementary Data 1, 2**).

**Fig. 2:**
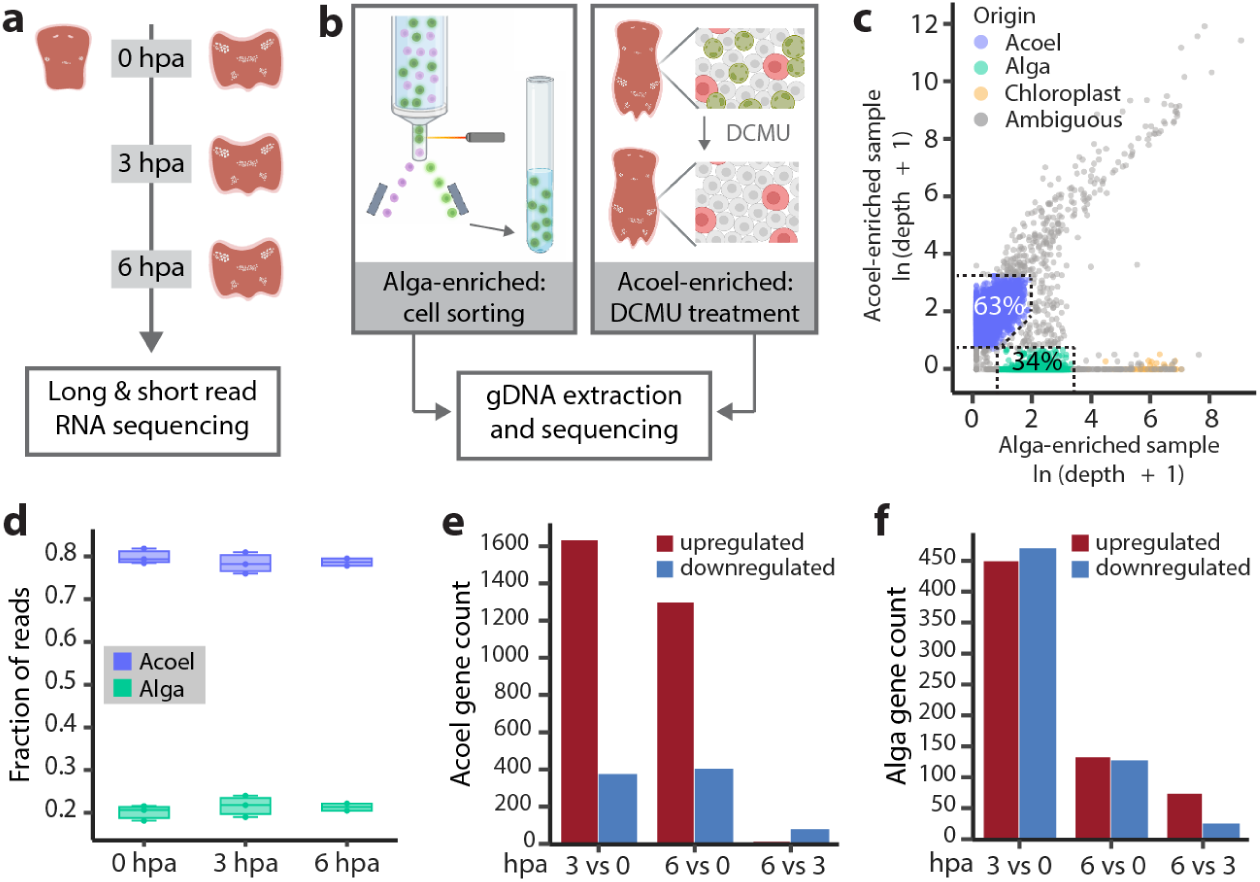
Assembly of reference transcriptomes enables the measurement of molecular wound responses in both acoel and algal cells. **a** Tails at 0, 3, and 6 hpa and heads at 0 hpa were collected for long and short read RNA sequencing to assemble reference transcriptomes *de novo* for both the acoel and the alga. Three replicates were obtained from 0 and 3 hpa tails, two from 6hpa, and one from the 0 hpa head, with each replicate containing 5 animals. **b** Experimental design for identifying the species origin of transcripts through sequencing the DNA extracted from acoel-enriched and algae-enriched samples. **c** Sequencing depth of each transcript in the acoel-enriched and algal-enriched samples. Dotted lines indicate the gates used to assign acoel and algal origin. The percentage of transcripts classified in each category is shown. Chloroplastic transcripts are annotated based on their high abundance in the algal-enriched sample and GO term predictions (cellular component GO term: 0009507 or 0009535, localized in the chloroplast). **d** Fraction of acoel and algal reads is consistent across samples and treatments, based on the Illumina short-read sequencing of tail samples specified in (**a**). **e,f** Number of DEGs in acoel (**e**) and algae (**f**) at different time points post amputation, (p-adj ≦ 0.05, log_2_FoldChange ≧ 1 or ≦ -1).

To separate the transcripts based on species, we separately sequenced DNA from acoels treated with DCMU to eliminate algae (acoel-enriched sample), and flow sorted algal cells (algal-enriched sample) (**Fig. 2b**, see methods). After aligning the reads to the transcriptome, we quantified the depth (the number of reads mapped to the transcript normalized by the transcript length) and coverage (the percentage of the transcript covered by sequencing reads) of each transcript in both samples (**Fig. 2c, Supplementary Fig. 5)**. Since the input for this experiment is genomic DNA (gDNA), transcripts encoded by the same genome should have similar sequencing depths and should be enriched in one species over the other. Based on this, we selected conservative gates to separate acoel and algal transcripts, which are also supported by the differential sequencing coverage and contrasting GC content of transcripts between the two species (**Supplementary Fig. 5**). With this classification, we recovered 13,313 acoel transcripts and 7,216 algal transcripts. The species origin of a small number of transcripts (662) remained ambiguous (**Fig. 2c**), which may be encoded by high copy number genes within the acoel or algal genomes, mitochondrial and chloroplast genomes, or from bacterial and viral contamination. Indeed, among the ambiguous transcripts with high depth in the algae-enriched sample, we annotated 54 chloroplastic transcripts based on the Gene Ontology (GO) term predictions (**Fig. 2c**) including two homologues of the large subunit of Rubisco and multiple subunits of PSI, PSII, and ATP synthase.

Finally, we annotated the transcriptomes through BLAST and Trinotate (**Supplementary Fig. 5, Supplementary Table 2**). The resulting transcriptomes have BUSCO scores comparable to other species of acoels and algae (**Supplementary Table 3**). The average mapping rates of Illumina sequencing reads to these transcriptomes were 94.6% for all our RNA sequencing (RNA-seq) experiments, allowing us to reliably compare gene expression across conditions.

To analyze gene expression in each symbiotic species, we normalized the fractional coverage of acoel and algal transcripts separately. While the read count ratio of acoel to algal transcripts remained consistent after injury (**Fig. 2d**), we identified hundreds of differentially expressed genes (DEGs) in both acoel and alga at 3 hpa compared to 0 hpa, and these responses were mostly conserved at 6 hpa (**Fig. 2e,f**). This result confirms our hypothesis that algae respond to injury with large transcriptional changes in a time frame matching the host molecular wound responses.

### Algal transcriptional responses to host injury concentrate in photosynthesis, chlorophyll biosynthesis, and carbon concentrating mechanism

To identify algal pathways that respond to host amputation, we conducted a GO term analysis on the differentially expressed agal transcripts. Multiple terms related to photosynthesis, as well as chlorophyll and carotenoid biosynthesis, were upregulated (false discovery rate, FDR ≦ 0.1) at 3 hpa (**Fig. 3a**). In contrast, genes related to cell division, cell motility, and carbohydrate metabolism were downregulated (FDR ≦ 0.1) in algae at the same time point, consistent with the observation that algae do not repopulate the wound site at these early regeneration stages.

**Fig. 3:**
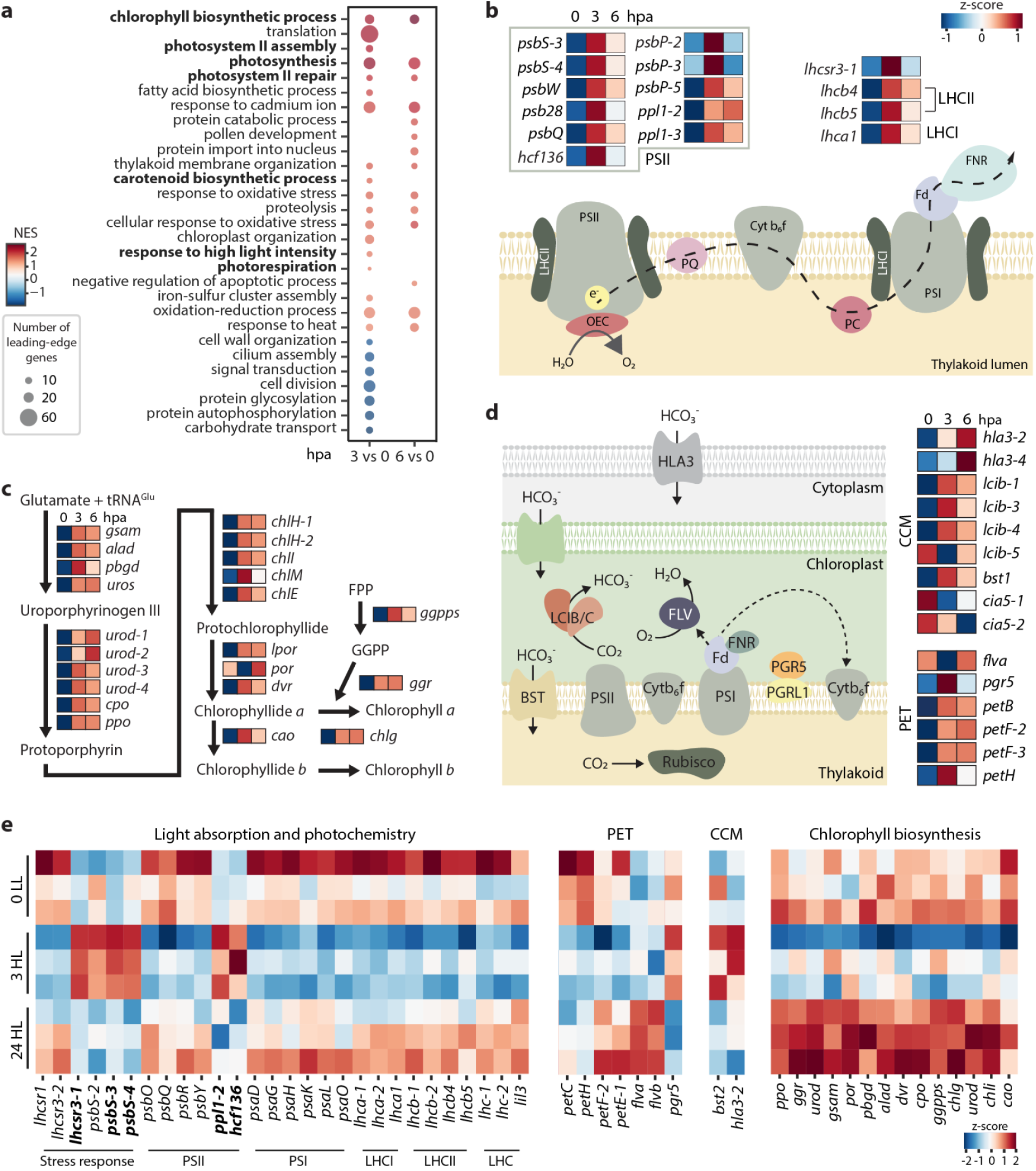
Host injury induces distinct molecular responses in algae. **a** GO term analysis of algal DEGs. Bolded GO terms are related to photosynthesis, chlorophyll and carotenoid biosynthesis pathways. **b-d** Diagram of the photosynthetic pathway (**b**), chlorophyll biosynthesis (**c**), carbon concentrating mechanisms (CCM) and alternative electron flow machinery (**d**), displaying DEGs (p-adj ≦ 0.05, log_2_FoldChange ≧ 1 or ≦ -1 respectively). Heatmaps report z-scores of average expression at 0, 3, and 6 hpa measured from three biological replicates, each containing 5 acoels. LHCII: light harvesting complex of PSII, LHCI: light harvesting complex of PSI, e^-^: electron, PQ: plastoquinone, Cyt b_6_f: cytochorme b_6_f, PC: plastocyanin, FPP: farnesyl pyrophosphate, GGPP: geranylgeranyl pyrophosphate. **e** Heatmaps showing normalized expression of genes associated with photosynthesis, alternative electron flows & CCM, and chlorophyll biosynthesis under low light (0 LL), after 3 hr exposure to high light (3 HL), and 24 hr exposure to high light (24 HL). Each row represents a biological replicate containing 5 acoels. Bolded transcripts are affected similarly at 3 HL and 3 hpa. PET: photosynthetic electron flow. CCM: carbon concentrating mechanism.

Examining specific genes supporting the GO-term enrichment, we found nineteen genes in the photosynthetic pathway collectively upregulated at 3 hpa (log_2_FoldChange ≧ 1, p_adj_ ≦ 0.05) (**Fig. 3b**). This list includes several components of PSII, including the accessory subunits *psb28* and *hcf136,* thought to be required for PSII *de novo* assembly and repair ^39,40^, and multiple homologs of *psbS*, reported to regulate light stress responses in other algae ^41^. We found similar trends in enzymes responsible for stabilizing the reaction center in the oxygen evolving complex (OEC) in plants and green algae, *psbP* and *psbQ* ^42^, and in components of the light harvesting complexes (LHC) of both PSI and PSII, which are essential for the capacity to absorb light energy ^43^ (**Fig. 3b**). Interestingly, the *light harvesting complex stress-related 3* (*lhcsr3*), involved in safe dissipation of excess light energy into heat in microalgae ^44^ is upregulated transiently at 3 hpa (**Fig. 3b**).

We also observed 21 genes involved in chlorophyll biosynthesis upregulated after host injury (**Fig. 3c**). Such upregulation of components of the photosynthetic complexes and chlorophyll biosynthesis has been implicated as part of a priming program that triggers a tolerance response in stress resistance in plants ^45–47^. The carotenoid biosynthesis pathway was also upregulated at 3 hpa. This pathway is known to produce pigments commonly used by a variety of phototrophic organisms for photoprotection ^48^, which implies that host injury induces a photoprotective or acclimation response in the algae.

During photosynthesis, electrons generated by photochemistry at PSII are transferred through the cytochrome b_6_f to PSI, creating high energy electrons used to reduce ferredoxin (Fd, *petF*) (**Fig. 3b**). The ferredoxin:NADP(H) oxidoreductase (FNR, *petH*) then generates reduced NADPH for CO_2_ fixation in the process known as linear electron flow. Both *petF* and *petH* were upregulated after injury (**Fig. 3d**). In addition, alternative electron flows using reduced Fd have been described including cyclic electron flow (CEF) and pseudo-cyclic electron flow (PCEF) ^49^. During CEF,electrons are transported around PSI via Fd and through cytochrome b_6_f, and this activity is regulated by *pgr5* and *pgrl1* ^50^. The *petB* subunit of cytochrome b_6_f, and *pgr5* are upregulated after injury (**Fig. 3d**). During PCEF, the electrons are used by flavodiiron proteins for the conversion of oxygen to water ^51^. After injury, we also observe an upregulation of one of the two flavodiiron genes (*flva*) at 3 hpa. These results are in contrast with the reduction of photosynthetic electron transport capacity upon injury as measured by a decrease in F_v_/F_m_ and Y(II) upon host injury, suggesting that algae cells compensate for loss of photosynthetic capacity with a higher transcription of these components.

In order to provide sufficient CO_2_ to the CO_2_-fixing enzyme Rubisco, microalgae have evolved CO_2_-concentrating mechanisms (CCM) that transport and concentrate inorganic carbon (CO_2_, HCO_3_ ^-^) inside the chloroplast for its assimilation by Rubisco ^49^. After amputation, three homologs of the low CO_2_-inducible proteins b (*lcib*) ^52,53^, were upregulated at both 3 and 6 hpa, along with bicarbonate transporters of the plasma membrane (*hla3*) ^54^ and thylakoid membrane (*bst1*) ^55^ (**Fig. 3d**). We also detected downregulation of the transcription factor *cia5*, which regulates CO_2_ responsive genes ^56^. These observations indicate that the inorganic assimilation pathway is modified upon amputation, potentially reflecting a decrease in inorganic carbon availability.

Since upregulation of genes involved in photoprotective mechanisms like *lhcsr* or *psbS* are recurrent in microalgae subject to a high light stress ^36^, we compared molecular responses to host injury and light stress. Upon high light stress, most genes involved in light absorption and photochemical reactions were downregulated acutely in response to high light exposure at 3 hr (labeled as “3 HL”) and returned to the baseline by 24 hr of high light exposure (“24 HL”) (**Fig. 3e**). This is in concordance with previous studies in plants and free-living microalgae under light stress ^36,38^. Only a handful of genes responded similarly at 3 hr after either light stress or amputation, including a subset of *psbS, psbP, lhcsr3-1,* and *hcf-136* homologs, which potentially represent generic stress responses as they are also induced by various stresses like CO_2_ limitation ^57^ and UV exposure ^36^ in other green microalgae.

We noted a similar difference in the chlorophyll biosynthesis pathway, with most upregulated genes at 3 hpa being downregulated in 3 HL samples, with some becoming upregulated in 24 HL (**Fig. 3e**). Genes involved in photosynthetic electron flow and CCM exhibited more complex patterns under high light, which did not overlap with injury-induced responses (**Fig. 3e**). These observations, combined with the photosynthetic efficiency measurements (**Fig. 1j-m**), suggest that both light and injury converge on affecting the algal photosynthesis, but through distinct sets of molecular factors.

### *runt* transcription factor is essential for acoel regeneration

To determine if algal injury responses are guided by the host regeneration program, we first focused on identifying regulators of acoel regeneration in order to perturb the host regeneration program and evaluate how it affects the algal responses. We found several acoel candidate regulators upregulated at 3 or 6 hpa (**Supplementary Fig. 6**). These include a set of conserved RNA binding proteins often associated with multipotentency in diverse organisms, such as homologs of *vasa, piwi,* and *argonaute 2* ^58^, along with conserved transcription factors (TFs), such as *egr, runt, fosl,* and *klf* homologs, some of which have been reported to play important roles in regeneration of other species ^28,29,59^. To further narrow down the list, we compared DEGs at 3 hpa between *C. longifissura* and the non-symbiotic acoel *Hofstenia miamia* that has been studied for its regeneration capabilities ^28^. We found only a handful of genes differentially expressed in both species at this time point (**Fig. 4a**). Of these shared DEGs, two are transcription factors: *runt* and *egr* (**Fig. 4a**), which we further analyzed in additional expression and functional studies.

**Fig. 4:**
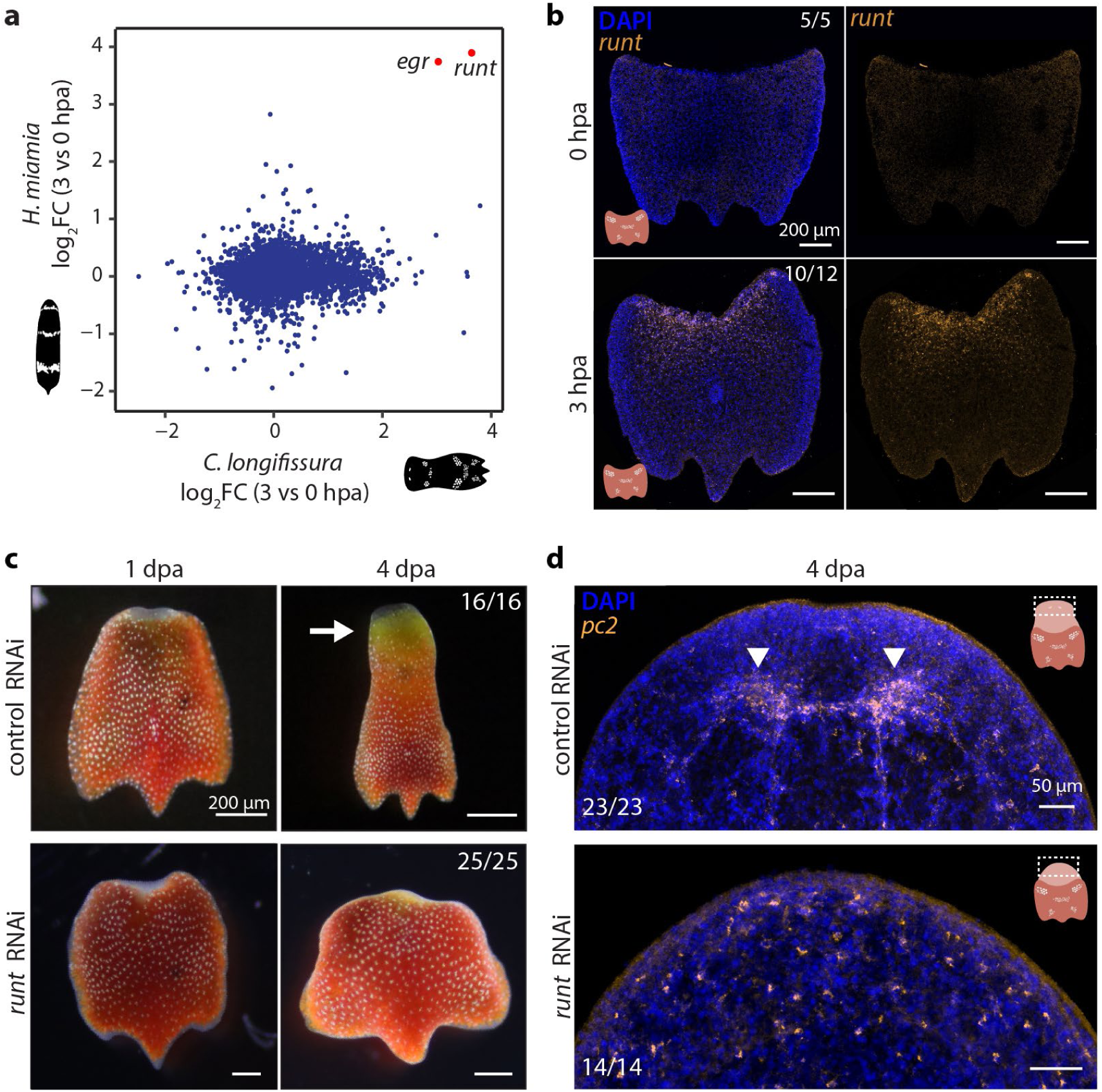
*runt* transcription factor is a conserved regulator of acoel regeneration. **a** Comparison of log_2_FC (fold change) in expression of orthologous genes at 3 vs 0 hpa between *C. longifissura* and *H. miamia*. One-to-one orthologs were identified using reciprocal BLAST. *H. miamia* data was obtained from Gehrke, *et* al., 2019^28^. **b** Expression of *runt* is evident only at 3 hpa at anterior wounds. Numbers represent animals in which the expression of *runt* is consistent with the figure out of the total number of animals examined. **c** Morphological comparison between *runt* and control RNAi treated animals at 1 and 4 dpa. While runt RNAi tail fragments can form a blastema at 1 dpa, they fail to regenerate head structures and eye spots, and have minimal tissue growth and shape changes. Numbers represent animals that showed regeneration phenotypes consistent with the images out of total animals examined. Arrow points at the regenerated head. **d** Expression of *pc2*, a neuronal marker, at 4 dpa in control and *runt* RNAi treated acoels. Neuronal ganglion is only present in control animals, whereas only distributed neurons are observed in *runt* RNAi treated animals. Numbers represent the animals that showed regeneration phenotypes consistent with the images out of total animals examined. Arrowheads point at the regenerated lobes of the neuronal ganglion in the control RNAi animal.

We validated the induction of *egr* and *runt* after injury in *C. longifissura* using in situ hybridization (**Fig. 4b**, **Supplementary Fig. 6**). While *egr* was activated in both anterior and posterior wounds, *runt* expression was specific to anterior wounds, clearly demonstrating the difference between the anterior and posterior regeneration programs. We proceeded to knock down *runt* and *egr* via RNAi. None of the *runt* RNAi tail fragments were able to regenerate a head, evidenced by the absence of the neural ganglion at 4 dpa along with other head structures (**Fig. 4c,d**), whereas only ∼15% of *runt* RNAi treated anterior fragments failed to regenerate a tail (**Supplementary Fig. 6**). Knockdown of *egr* led to regeneration deficiencies at a lower penetrance -- only a third of either heads or tails failed to regenerate (**Supplementary Fig. 6**). The highly reproducible anterior regeneration phenotype of *runt* RNAi was used to evaluate whether and how the algal response to injury depends on the host.

### *runt* RNAi alters algal response to host injury

To determine whether the algal transcriptional responses to host injury depend on the regeneration program controlled by *runt*, we performed RNA-seq on animals at 0 and 3 hpa after *runt* RNAi (**Supplementary Fig. 7**). Strikingly, this identified a large number of DEGs not only in the acoel, but also in algae at both 0 and 3 hpa, suggesting that *runt* influences the transcription in both partners, likely through intermediate signaling molecules or other factors in the symbiont.

We noticed that RNAi treatment significantly reduced *runt* expression, but did not entirely eliminate it after host injury (**Fig. 5a**). This enabled us to calculate the correlation between *runt* expression and the expression of all other genes, providing an alternative analysis to identify genes modulated by *runt* RNAi in both acoel and algae.

**Fig. 5:**
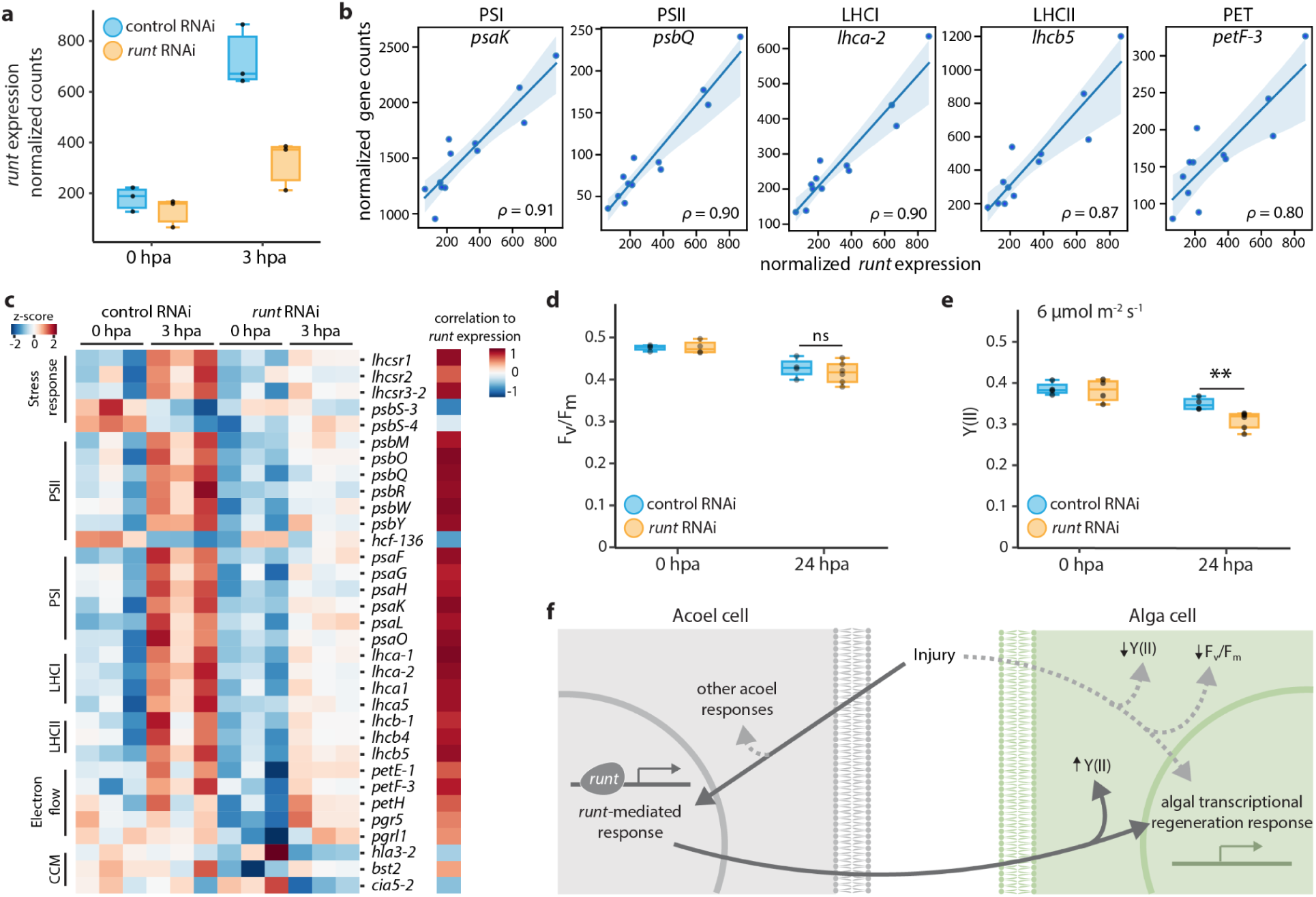
*runt* affects transcription of photosynthetic related genes and photosynthetic efficiency. **a** Expression of *runt* at 0 and 3 hpa in control and *runt* RNAi acoels. **b** Normalized expression of example photosynthesis genes vs *runt* normalized counts. Each dot represents a sample from the RNA-seq experiment. Replicates are shown as dots, line represents the linear regression, and shaded region is the confidence interval of the regression. ρ: Spearman correlation. The examples are chosen to represent components of PSI, PSII, LHCI, LHCII, and PET, respectively. **c** Left: heatmap reporting z-scores from triplicates of selected genes involved in the light reactions of photosynthesis, alternative electron flows, and CCM that are differentially expressed at 3 hpa in *runt* RNAi vs control RNAi, or highly correlated with *runt* expression. Right: correlation of the given gene with *runt* expression. **d** F_v_/F_m_ measured at 0 and 24 hpa in *runt* RNAi and control tails. There is a similar decrease in both *runt* RNAi and controls after 24 hpa. **e** Y(II) of *runt* RNAi and control tails at 0 and 24 hpa with 6 µmol photon m^-2^ s^-1^ of actinic background light. Both *runt* RNAi and controls show a decrease in Y(II) at 24 hpa compared to their counterparts at 0 hpa. Y(II) of *runt* RNAi at 24 hpa compared to control RNAi is significantly reduced (**pval ≦ 0.05, ns, not significant, t-test). For (**e**) and (**f**) each treatment consisted of four biological replicates measured in parallel, and each replicate contained twenty tails. **f** Schematic of proposed *runt*-mediated coordination between acoel and algal wound responses. Dark gray arrows indicate responses dependent on *runt* activation. Activation of algal responses that are dependent on *runt* should be regulated indirectly through intercellular communication. The light gray, dashed arrows represent alternative pathways that may be activated during regeneration and influence algal responses in a *runt*-independent manner.

We identified algal genes that may be activated by the *runt-*mediated injury response by selecting genes that are either significantly downregulated in *runt* RNAi animals at 3 hpa (p-adj ≦ 0.1, log_2_FC ≧ 0.8) or in strong positive correlation with *runt* expression (***p*** ≧ 0.8). Of the 91 genes selected, twenty are related to light harvesting and photochemical reactions, of which eight are LHC proteins and twelve are subunits of PSI and PSII (**Fig. 5b,c**). Many of these genes continued to respond to injury in the *runt* knockdowns, but their expression was also dampened. Importantly, genes involved in photosynthetic electron transport including the cytochrome b_6_f subunit, the plastocyanin *petE*, the ferredoxin *petF*, the FNR *petH*, and multiple PSI and PSII subunits were correlated with *runt* expression, indicating that *runt* expression may contribute to sustain the photosynthetic electron transport in algae after host injury. To test whether physiological changes also depend on *runt*, we measured the photosynthetic efficiency after *runt* RNAi. While the knockdown of *runt* did not affect F_v_/F_m_ at either 0 or 24 hpa (**Fig. 5d**), Y(II) was significantly reduced at 24 hpa compared to the control (**Fig. 5e**, **Supplementary Fig. 7**). These results imply that, while other injury-induced factors may be responsible for the reduction of the algal maximal photosynthetic capacity, likely via degradation of PSII, the *runt-*dependent responses contribute to sustaining the electron transport activity after injury.

We also observed multiple transporters affected by the *runt-*mediated response in both acoel and algae. In the algae we found seven transporters dependent on *runt* expression, including a bicarbonate transporter (*hla3-2*), two *zinc-iron permeases* (*zip3* and *zip12*), two sodium:solute transporters (*slc5a7* and *slc6a13*), and a *urea active transporter* (*dur3*) (**Supplementary Fig. 7**). In the acoels, we found fourteen transporters affected by *runt* knockdown. Two of these, *slco4c1* and *vha,* are downregulated by *runt* expression in response to injury and could be regulating the acidification of the extracellular environment ^19,20,60^. A glutamate transporter, *eaat1*, is also downregulated by *runt* and could be affecting the nitrogen cycling between the symbionts. Multiple solute carriers were upregulated by *runt*, including *slc18b1, slc17a5, slc6a18, slc6a8,* and *slc6a5,* which may modify the nutrient exchange with the algae during regeneration ^61^ (**Supplementary Fig. 7**). This suggests that *runt* is likely involved in mediating the acoel-alga communication during regeneration, by tuning the exchange of metabolites and small molecules between the two organisms.

## Discussion

In this study, we establish the photosymbiotic acoel *C. longifissura* as a system for dissecting the molecular interactions between species during whole body regeneration, using a suite of sequencing approaches and functional genomic analysis. We demonstrate that in addition to the animal’s early wound response, endosymbiotic algae respond physiologically and transcriptionally to host injury. The symbiotic algae reduced their photosynthetic efficiency drastically and upregulated genes related to light harvesting and photochemistry, chlorophyll biosynthesis, and carbon concentrating mechanisms. Intriguingly, upregulation of photosynthetic and pigment related genes has been described in stress-tolerant plants and tolerance primed plants ^45–47^. In contrast, light stress led to the downregulation of an overlapping set of genes while also causing a decrease in photosynthetic efficiency. These observations suggest that, unlike responses to light stress that aims to contain photo-damage, the injury-induced transcriptional responses may compensate for the loss in photosynthesis due to host injury.

Another important result from this study is that one of the acoel’s regeneration-responsive pathways, which is controlled by the conserved transcription factor *runt*, contributes to the algal responses to host amputation, albeit indirectly (**Fig. 5f**). *runt* knockdown affected Y(II) but not F_v_/F_m_, suggesting that the changes caused by *runt* activation impact the efficiency of photosynthetic electron flow rather than the availability of functional PSII. Furthermore, the injury-induced upregulation of genes involved in the photosynthetic electron flow also depended on *runt* expression. The dampened transcriptional response after *runt* knockdown could be translated into a slower repair of damaged components of the electron transport chain.

Comparing our results with previous studies in another non-symbiotic acoel, *H. miami*a ^28^, it is plausible that *runt* is a conserved master regulator of acoel regeneration, as it is activated after injury and essential for regeneration in both acoels, suggesting that *runt* activation has been co-opted to activate algal responses in *C. longifissura*. However, we noticed that there is minimal overlap in wound responses between the two acoels, indicating that the endosymbiotic algae may have modified the regeneration response pathways in *C. longifissura*, as endosymbiosis in acoels is a derived trait whereas regeneration is shared by multiple acoel clades.

As an animal transcription factor, it is unlikely that *runt* could directly control transcription in algae. While it remains unclear how the host and the symbiont cells communicate, we noted that several transporters appear to be regulated by *runt* in both animals and algae (**Supplementary Fig. 7**), implying that the exchange of metabolites or signaling molecules may be involved in this process. Indeed, nutrients, nitrogen-based compounds, and organic and inorganic carbon exchange occur regularly between endosymbiotic partners during homeostasis and are modified during stress in other organisms ^17,62–65^. This exchange may lead to a modified pathway that allows for coordinated regeneration responses in both the host and symbiont.

Two candidate communication pathways between host and symbionts are regulated by nitrogen and carbon metabolites. The algal urea transporter *dur3* is downregulated in parallel to the acoel’s *glutamate synthase* (*gogat*), *glutamate transporter* (*eaat1*), and *ammonium transporter 1* (*amt-1*) after amputation in *runt* RNAi animals (**Supplementary Fig. 7**). It has been reported that nitrogen availability is important for photosynthesis and proliferation in algae and can serve as a mechanism for the host to regulate algal endosymbiont proliferation in other systems ^17,66^. Similarly, carbon exchange could be playing a role during regeneration. The availability of inorganic carbon is required for algal carbon fixation while the heterotrophic host requires organic carbon compounds for survival ^19,20,49,62^. Recent studies suggest that in addition to the algal CCM, animal hosts also concentrate carbon for their algal symbionts ^19,20^ with V-type H^+^-ATPase (VHA) acidifying the microenvironment and promoting the conversion of HCO_3_ ^-^ into available CO_2_ for the algae. In *C. longifissura*, *vha* and *slco4c1,* an organic anion transporter that may use HCO_3_ ^-^ as a counterion ^60^, were negatively correlated to *runt* expression (**Supplementary Fig. 7**). Conversely, on the algal side, the CCM master regulator *cia5* and the bicarbonate transporter *hla3-2* were also negatively correlated to *runt* (**Fig. 5c**). By regulating carbon and nitrogen availability, these transporters could be an acoel-algal communication strategy to trigger the algal response to injury.

Algae may also recognize host injury without the need of direct communication, but through sensing common regeneration-induced physiological changes, such as reactive oxygen species (ROS) and ascorbic acid, an antioxidant that scavenges ROS ^67^. While elevated ROS levels have been demonstrated during tissue regeneration in several animals ^68,69^, in photosynthetic organisms ROS is inducing transcriptomic responses that regulate most of the genes involved in light harvesting and photosynthetic electron flow ^70,71^. In *C. longifissura*, *runt* activation leads to an increase in the expression of *gulonolactone oxidase* (*gulo*), which is responsible for ascorbic acid biosynthesis in animals ^72^. Changes in both ROS and ascorbic acid concentrations may be recognized by the algae and activate their responses.

It is worth mentioning that other factors, independent of *runt*, should also contribute to the algal responses, as algae continued to have some physiological and transcriptional responses to host injury after *runt* RNAi. These may include microenvironment changes caused by wounding, such as fluctuations in osmolarity, pH, and even light penetration, as well as other acoel molecular wound responses that are controlled by separate pathways.

Finally, although our experiments suggest that algal photosynthesis is not required for acoel regeneration, other aspects of the algal biology may be required during this process. The previously discussed metabolic exchange mechanisms could be an avenue for future studies. These exchanges may also affect asexual reproduction, a trait that is more common in the Convolutidae family, with the *Convolutriloba* genus presenting multiple unique fission and budding strategies ^25^. In other *Convolutriloba* species, an accumulation of algal symbionts has been observed prior to asexual division ^24,73^. If and how the endosymbiotic state may have led to the evolution of this trait is another intriguing question. Differences between embryonic development and the regeneration program, as well as comparison between regeneration of other symbiotic and non-symbiotic acoels, may determine if and what modifications occur in the presence of algae.

## Materials and methods

### Animal maintenance

Acoels were cultured in a 16 gallon Coralife Biocube Aquarium in artificial seawater (ASW, 34 ppt, Instant Ocean) at 26 °C with a 14h:10h light:dark cycle. Acoels were fed freshly hatched *Artemia* shrimp every 1-3 days *ad libitum* in the tank. Animals used in experiments were kept in individual wells in an incubator at 26 °C with a 14:10h light:dark cycle, with filtered ASW. These animals were fed and their water was changed every 1-3 days. The light intensity to which the animals were exposed was modulated for individual experiments between 50 and 150 µmol photon m^-2^ s^-1^.

### Regeneration experiments

Animals were kept individually or in groups smaller than 10 individuals in 12-well or 24-well plates in an incubator (Danby Fresh 1.7 cu. ft. Herb Grower, cat. #DFG17A1B). Pictures of animals were taken at the same time every day, over four days after amputation, using a stereo microscope (Zeiss Stemi 508) and an inverted fluorescence microscope (Zeiss Axio Observer Z1) to evaluate acoel and algae growth. For dark treatment, a chamber was used to keep acoels in the dark at 26 °C. Animals were not exposed to light before imaging. Individual animals were only imaged once. For long dark treatment, acoels were kept in the dark for two weeks prior to amputation and fed every other day. All treatments were replicated at least three times with at least three animals per replicate.

### DCMU treatment

Acoels were treated with 20 μM DCMU in ASW, by diluting the stock solution of 20 mM DCMU in ethanol. Vehicle controls were treated with ethanol (1:1000). After 24 hr, acoels were rinsed multiple times and maintained in fresh ASW. Effects of the DCMU treatment were confirmed with a Dual-PAM system, showing blockage of PSII (**Supplementary Fig. 3**). For long-term DCMU treatments, animals were treated with DCMU twice, three days apart. Algae were completely removed from the acoel two weeks after the two treatments. The removal of algae was validated through the red fluorescence of the algae (∼633 nm).

### Photosynthesis measurements

To measure chlorophyll fluorescence in the algae within the animal, we created a chamber for containing acoels on a Dual-PAM 100 (Walz) pulse amplitude modulation (PAM) fluorimeter using a small plastic Petri dish and a 2 mm thick silicone container (**Supplementary Fig. 2**). We carved a 5 × 6 mm space to contain the animals. On the edges of the acoel chamber we increased the height to 4 mm so that the diode of the emitter would not touch the water. This device was then mounted on a PDMS base that fit the detector diode in order to keep the sample stable during measurements. The Dual-PAM system was used in a vertical configuration (**Supplementary Fig. 2**). We used 20 tail fragments for all experiments.

After mounting the acoel chamber, chlorophyll fluorescence measurements were performed. Detection pulses (10 µmol photon m^-2^ s^-1^ blue light) were applied at a 100-Hz frequency. Basal fluorescence (F_o_) was measured after a 20 min dark adaptation prior to the first saturating flash. Red saturating flashes (6,000 μmol photon m^-2^ s^-1^, 600 ms) were delivered to measure F_m_ (in the dark) and F_m_^’^ (in the light). PSII maximum yields (F_v_/F_m_) were calculated as (F_m_-F_o_)/F_m_. The operating quantum yield of PSII (Y(II)) was calculated upon actinic illumination as (F_m_^’^ - F_o_)/F_m_. For light curves, animals were exposed to increasing light intensities, and for each light intensity, acoels were acclimated to the light for 2 min prior to a measurement of Y(II) in order to quantify steady state photosynthetic rates.

### RNA extractions

Three replicates each containing five animals were used for RNA extraction for each condition. We rinsed the animals in ASW twice and then removed as much ASW as possible. 300 μl of Trizol was added, followed by a 2-3 min incubation at room temperature and vigorous vortexing to dissociate the tissue. Samples were then flash frozen on dry ice and kept at -80 °C until extraction so that all samples for each RNA-seq experiment could be processed in parallel. On the day of the extraction, samples were thawed on ice and 700 μl of fresh Trizol was added. After brief vortexing, 200 μl of chloroform was added and the sample was shaken vigorously for 15 s followed by a 2 min incubation. Samples were then centrifuged (16,000 g, 15 min at 4 °C) and the aqueous phase was carefully transferred into new tubes and processed with the Direct-zol RNA Purification Kit (Zymo Research, cat. #R2051) following the manufacturer’s instructions, which includes a DNAse treatment step. RNA concentration and quality was quantified on a Bionanalyzer.

### RNA-seq library preparation

50 ng of input RNA was reverse transcribed (RT) using the Smartscribe Reverse Transcriptase (Clonetech). Full-length cDNA was generated using a modified Smart-seq2 protocol ^74^. During the RT reaction a template-switch oligo (TSO) and a custom oligo(dT) primer containing a UMI and sample barcode were supplemented to enrich mRNA (**Supplementary Table 4**). The RT reaction was performed in 10 μL reactions and incubated at 42 °C for 1 hr. After first strand cDNA synthesis, 1 μL of 1:10 dilutions of RNAse A (Thermofisher) and Lambda Exonuclease (NEB) were added and incubated at 37 °C for 30 min. Following the incubation, an amplification step was performed using KAPA Hifi ReadyMix 2X (KapaBiosystems) containing 1 μL of ISPCR primer (10 μM) in 25 μL reactions. Samples were incubated at 95 °C for 3 min, followed by 12 PCR cycles of (98 °C for 20 s, 62 °C for 15 s, and 72 °C for 4 min), with a final extension at 72 °C for 5 min. Libraries were purified using a ratio of 1:0.85 sample to bead ratio using Agencourt AMPure XP SPRI beads (Beckman Coulter). The final products were quantified on a D5000 Tapestation (Agilent) or a Bioanalyzer using the High Sensitivity DNA kit (Agilent, cat. #5067-4626).

To generate input cDNA (>1 μg) for producing Oxford Nanopore Technologies (ONT) and PacBio libraries, which were used for the transcriptome assembly, samples were pooled together equally with 20 ng of cDNA each and amplified through an additional 6 PCR cycles. The entire pool was purified using Agencourt AMPure XP SPRI beads at a 1:0.85 ratio and eluted in 50 μl yielding a final library concentration of ∼115 ng/μL.

For the ONT library preparation, ∼1-2 μg of the final full-length cDNA product was prepared using the ligation based ONT method with the SQK-LSK109 kit, according to the ONT instructions with minor modifications. First, the End Repair and A-tailing reaction was extended for 30 min each at 20 °C and 65 °C instead of 5 min each. Second, the ligation reaction time was also extended to 30 min at room temperature instead of 10 min per protocol. Two runs were performed using the MinION device using the 48 hr sequencing protocol in the MinION 9.4.1 chemistry flowcells. All bases were called with the high-accuracy GPU accelerated model of Guppy v3.5.2.

For the PacBio library preparation, a SMRT library was prepared with 1 μg of the full-length cDNA product using the Sequel II binding kit 2.0. Reads were processed through the Circular Consensus Sequencing (CCS) pipeline using Smartlink to generate high quality reads. Each read from the CCS output was generated using parameters of “min pass = 1” and “min accuracy = 0.85”.

For Illumina sequecing, 4-10 ng of the full-length cDNA was used as input for preparing Nextera XT (Illumina, cat. # FC-131-1024) libraries following the manufacturer’s recommendations. The input cDNA was indexed using a tagmentation reaction and then incubated at 72 °C for 3 min to heat inactivate the enzyme. The indexed cDNA libraries were amplified with 12 PCR cycles of (95 °C for 10 s, 55 °C for 30 s, and 72 °C for 30 s), with a final extension at 72 °C for 5 min. Some amplified libraries were size selected for 300 - 800 bp on a 2% EX E-gel (Thermofisher) and purified using QIAquick gel extraction kit (Qiagen). Libraries were pooled at equal concentrations and sequenced either on a NextSeq 500 using High Output runs, or on a NovaSeq 6000.

### Transcriptome assembly and annotation

We filtered Illumina reads using Trimmomatic (v 0.39) with the following parameters “LEADING:3 TRAILING:3 SLIDINGWINDOW:4:15 MINLEN:36”, and Nanopore and Pacbio reads using NanoFilt (v 2.7.1) with the following parameters “-q 10 -l 150 --headcrop 75 -- tailcrop 75”. We assembled an initial transcriptome draft with RATTLE assembler ^75^ using the filtered Nanopore reads. We then aligned filtered Illumina and Pacbio reads to the initial draft using Minimap2 (v 2.17-r941) to perform base polishing using Pilon (v 1.23) ^76^. To remove chimeric transcripts or regions with poor read support, we performed a coverage scan with a rolling window size of 10 bp to trim both ends of the transcript that have less than 33% of the maximum alignment coverage. After trimming, transcripts that have lengths shorter than 300 bp were removed. We then re-mapped the Illumina reads to the initial draft using Salmon quant (v 1.3.0) and clustered similar transcripts with Grouper (https://github.com/COMBINE-lab/grouper) to remove duplicate transcripts. Illumina and Pacbio reads were then re-mapped against the clustered transcriptome using Minimap2 for a second round of base polishing, yielding the final version of the transcriptome that we used for the remainder of our work.

We predicted the open reading frame (ORF) for each transcript using TransDecoder (v 5.5.0) (https://github.com/TransDecoder/TransDecoder) with default parameters, keeping only the predicted ORF with the longest length. We performed functional annotation using Trinotate (v 3.2.1). To identify putative homologs, we performed blastx on the transcriptome against the NCBI “refseq_protein” database. To evaluate the completeness of our transcriptomes, we performed BUSCO (v 4.0.5) analysis.

### Determining the species origin of transcripts using DNA sequencing

We treated a large cohort of acoels with two rounds of DCMU, each lasting 24 hr (see **DCMU treatment**), in order to remove algal cells and obtain an acoel-enriched sample. After two weeks of incubation, we confirmed the absence of algal cells through fluorescence imaging and selected animals with none or rare algal cells. We then proceeded to flash freeze these animals in liquid nitrogen and stored them at -80 °C.

The algal-enriched sample was collected on a cell sorter (Sony SH800S) based on algal autofluorescence (**Supplementary Fig. 5**). Acoels were dissociated on ice in the dissociation media (3.3x calcium magnesium free PBS, 2% FBS, 20 mM HEPES) by gently pipetting for 10 min until solution was homogenized. The solution was then filtered through a 40 μm strainer to remove debris and placed on ice. Before sorting, the solution was filtered again through a 35 μm strainer and gently mixed. Sorted algal cells were centrifuged and washed twice with ASW before gDNA extraction.

gDNA was extracted using a high-molecular weight DNA isolation protocol ^77^ with some modifications. We used 400 μl of GTC buffer with a 30 min incubation for tissue dissociation, then added 200 μl of distilled water and 400 μl of phenol/chloroform, and mixed by inversion. We then centrifuged at 12,000 g for 15 min at 4 °C and transferred the upper aqueous phase into a new Phase Lock gel tube. 500 μL of chloroform were added, and the solution was mixed by inversion before centrifugation at 12,000 g for 10 min at 4 °C. The upper aqueous phase was again transferred to a new Phase Lock gel tube. We then added 200 μL of 5 M NaCl, mixed by inversion before incubating for 10 min on ice, and centrifuged for 10 min. The upper aqueous phase was then transferred to a DNA LoBind tube, 600 μl of cold isopropanol were added, and samples were stored at 4 °C overnight. Samples were centrifuged for 2 hr at 4,000 g at 4 °C. The pellet was washed with 1 mL of 70% ethanol and centrifuged at 4,000 g for 10 min at 4 °C. The pellet was resuspended in 50 μL of TE buffer (10 mM Tris-HCl, 1 mM EDTA, pH 8.0). We quantified gDNA concentration on a Qubit. Libraries were prepared with the Illumina DNA Prep kit (cat. #20018704) following the manufacturer’s instructions. 72 ng and 420 ng were used as input for the alga-enriched, and the acoel-enriched samples, respectively. Reads were aligned to the transcriptome using Minimap2 (v 2.17-r941) with preset parameters for genomic short-read. We selected only properly paired mapped read counts for the analysis. Transcript depth and coverage was calculated with samtools (v 1.13, https://github.com/samtools/samtools).

### Differential gene expression analysis

Short reads were aligned with Minimap2 (v 2.17-r941), quantified with HTSeq, and differential gene expression analysis was performed using DESEQ2 (v 1.38.2), separately for the algal and acoel transcripts. For algal genes, the putative chloroplast genes were not included since our library preparation protocols enriched for polyadenylated transcripts and polyadenylation in chloroplast genes targets them for degradation instead of transcription ^78^. GO term analysis was performed using the GSEApy package (v 1.0.4) ^79^.

### BrdU staining

Acoels were exposed to 0.1 mg/ml BrdU for 2 hr, washed, and amputated. At 0 or 24 hpa, animals were relaxed in 3.4% MgCl_2_ for 5 min and then immediately incubated in 4% formaldehyde (FA) in ASW for 1.5 hr. Samples were washed with PBS and then denatured in HCl (3:1 in distilled water) at 37 °C for 30 min. Samples were washed with PBSTx (PBS supplemented with 0.1% Triton-X), and blocked in 10% goat serum in PBSTx (blocking solution) for 1 hr, followed by an incubation with mouse anti-BrdU monoclonal antibody (Sigma cat. #B2531, 1:100 dilution in blocking solution) overnight at 4 °C. Samples were then washed multiple times with PBSTx before being incubated in FITC-conjugated goat anti-mouse secondary antibody (Sigma cat. #A6667, 1:200) for 2-5 hr at room temperature. Finally, samples were mounted in 75% glycerol for imaging.

### In situ hybridization

Animals were rinsed with filtered ASW, relaxed briefly in 3.4% MgCl_2_, and fixed with 4% FA in ASW for 1.5 hr. Samples were then washed twice with PBSTw (PBS + 0.1% Tween-20) for 5 min each, followed by two quick rinses with 100% methanol. Fixed samples were stored in 100% methanol in -20 °C.

Riboprobes for in situ hybridization were synthesized using the oligonucleotide primers listed in **Supplementary Table 4** to clone the DNA fragment of interest into vector pJC53.2 (Addgene Plasmid ID: 26536), followed by riboprobe synthesis previously described ^80^.

Fixed animals were bleached in 6% H_2_O_2_ in methanol for 1 hr, washed with 100% methanol, rehydrated with 50% methanol:PBSTw, followed by two PBSTw washes. Then, they were incubated in Proteinase K solution (2 μg/mL supplemented with 0.1% SDS in PBSTw) for 10 min without shaking and immediately post-fixed with 4% FA in PBSTw for 1 hr. After two PBSTw washes, and one wash in 50% pre-hybridization buffer, samples were incubated in pre-hybridization buffer (prehyb, 50% deionized Formamide, 0.1% Tween-20, 5x SSC, 1% SDS in DEPC-treated water) at 56 °C in a hybridization oven for 2.5 hr. Hybridization proceeded overnight at 56 °C in hybridization buffer (50% deionized Formamide, 0.1% Tween-20, 5x SSC,1% SDS, 0.1 mg/mL yeast RNA, 0.1 mg/mL salmon sperm DNA, 62.5 μg/mL of heparin, in DEPC-treated water; riboprobes diluted at 1:1000). Samples were sequentially washed in prehyb, 50% prehyb, 2x SSC supplemented with 0.1% Tween-20, and 0.2x SSC with 0.1% Tween-20 for 20-30 min each at 56 °C. Samples were brought back to room temperature during the last wash, and then washed twice with MABTx (11.6 g/L maleic acid, 8.8 g/L NaCl, pH 7.5, 0.1% TritonX). Blocking was performed in 10% horse serum in MABTx for 1 hr, followed by incubation in an antibody solution overnight. For fluorescence in situ hybridization (FISH), we used anti-dig-POD (Roche, cat. # 11207750910, 1:1000 in blocking buffer) and for whole mount in situ hybridization (WISH) we used anti-digoxigenin-AP antibody (Roche cat. # 11093274910, 1:4000 in blocking buffer).

For WISH, we washed the samples ten times in PBSTx supplemented with 0.1% BSA (PBTx), each for 20 min. Samples were incubated in AP buffer (0.1 M Tris, 0.1M NaCl, 0.05 M MgCl_2_, 0.5% Tween-20), and then development buffer (4.5 μL/mL NBT, 3.5 μL/mL BCIP in AP buffer). The reaction was stopped with PBTx, fixed in 4% FA for 10 min, rinsed again in PBTx, and washed in 100% ethanol until color turned dark blue. Samples were rinsed in PBTx and mounted for imaging in 80% glycerol supplemented with 10 mM Tris and 1 mM EDTA, pH 7.5.

For FISH, we washed the samples five times in MABTx and five times in PBTx, each for 20 min. We then incubated the samples in Tyramide Buffer (2 M NaCl + 0.01 M Boric Acid, pH 8.5) for 10 min, followed by a 10 min incubation in the Development Buffer (20 μg/mL IPBA, 0.003% H_2_O_2_, 20 μg/mL TAMRA in Tyramide Buffer) in the dark. After washes in PBTx, samples were mounted in scale solution (30% glycerol, 0.1% TritonX, 4 M Urea, 2 mg/mL sodium ascorbate, in PBS) for imaging. Samples were imaged on a Zeiss LSM 800 confocal microscope. Confocal sections with optimal z spacing based on the Zen software were recorded to image the entire thickness of the acoel and maximum intensity projections were generated.

For HCR, samples were instead incubated with probe hybridization buffer (Molecular Instruments) for 30 min at 37 °C, before the hybridization buffer (probe hybridization buffer with 4 pM of the *runt* oligo pool) was added. The *runt* oligo pools were designed using the probe generator from the Ozoplat Lab ^81^ (**Supplementary Table 4**). Samples were incubated for 12 hr, and washed four times with probe wash buffer (Molecular Instruments) at 37 °C and five times at room temperature with 5x SSCT (5x SSC, 0.1% Tween-20). Samples were then incubated in Amplification Buffer (Molecular Instruments) for 30 min and incubated in a hairpin solution (30 pMl of each hairpin, heated to 95 °C and snap cooled in Amplification Buffer). Samples were washed multiple times with 5x SSCT and mounted in scale solution (30% glycerol, 0.1% Triton X-100, 2 mg/mL sodium ascorbate, 4 M urea in 1x PBS) ^82^.

### RNAi-mediated gene knockdown

Genes of interest were identified from our newly assembled transcriptome and primers were designed for the predicted ORFs (**Supplementary Table 4**). Double stranded RNA (dsRNA) was synthesized using pJC53.2 plasmids (Addgene Plasmid ID: 26536), as previously described ^80^, and resuspended in 50 μL MilliQ water. RNAi was performed through microinjection into the gut using a XenoWorks Digital Microinjector. Needles were pulled with a vertical micropipette puller (Sutter Instruments Model P-30, with settings heat: 750 and pull: 750). The dsRNA was diluted 1:1 with ASW and mixed with food coloring dyes for visual confirmation of successful delivery. *runt* RNAi injections were done every 2 days over a week (3 injections total). *egr* RNAi injections were done every 2-3 days for three weeks (9 injections). Animals were fed brine shrimp before injections to ensure the formation of the acoel gut, and water was changed every couple of days. As the negative control, acoels were injected with the *ccdB* and *camR* insert from the pJC53.2 plasmid. Control RNAi was administered with the same frequency and for the same time span as *runt* or *egr* respectively.

### Code and data availability

All sequencing data will be available through NCBI. The transcriptome assembly and annotation pipeline will be available at www.github.com/xuesoso/acoel_reference_assembly.

## Acknowledgments

We thank P Bump for help with designing the oligo pools used for the *runt* HCR experiment, J Gibson for help with the gDNA extraction protocol, and all Wang group members for discussion. DNS is a BioX Bowes Fellow, ES and YX are Stanford Interdisciplinary Graduate Fellows. BW is a Beckman Young Investigator. ABurlacot is supported by the Carnegie Institution for Science. This work is supported by a NIH grant 1R35GM138061 to BW.

## Author contributions

Conceptualization: DNS, YX, JS, and BW; Methodology: DNS, YX, AByrne, DL, ABurlacot, and JS; Investigation: DNS, YX, and ES; Formal analysis: DNS, YX, and ES; Validation: DNS; Writing: DNS and BW with feedback from YX, ABurlacot, and JS; Funding acquisition: BW; Supervision: SD, SRQ, ABurlacot, JS, and BW.

## Supplementary Figures & legends

**Supplementary Figure 1:**
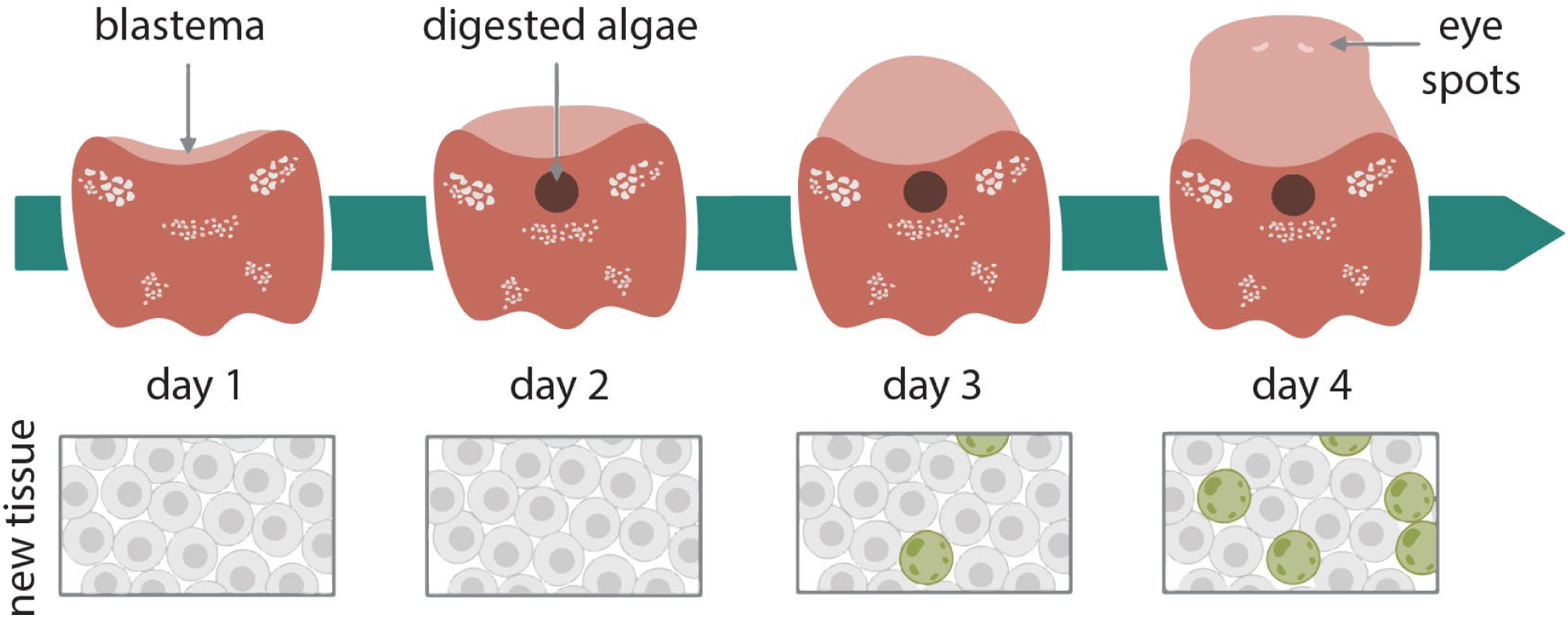
Schematic showing the process of anterior regeneration from *C. longifissura* tail fragments. Animals are bisected at the posterior pair of white concrement granules, which coincides with the fission plane during asexual reproduction. Lighter color depicts the new tissue, with darker color representing the pre-existing tissue. A black spot often forms at 2 dpa, which has been suggested to be the accumulation of digested algae^24^. Lower panels: algal population in the new tissue. Algae (green) begin to populate the new tissue at 3 dpa and completely cover it by 4 dpa.

**Supplementary Figure 2:**
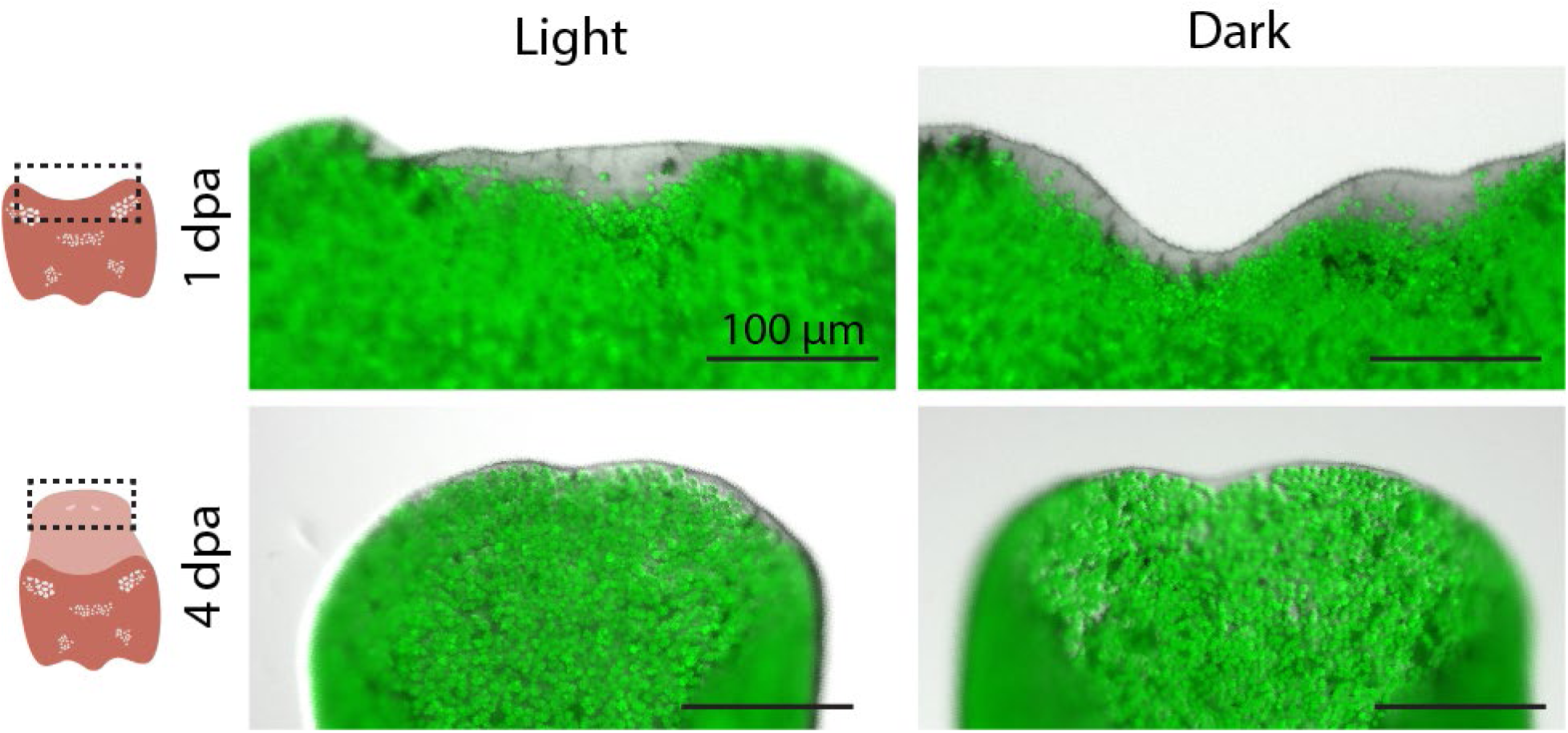
Dark treatment does not affect acoel regeneration or algal repopulation of the new tissue. Anterior regeneration in the presence or absence of light during regeneration at 1 and 4 dpa. Regeneration appears to be consistent and happens at a similar pace between conditions. Algal cells (green) are imaged through autofluorescence at 647 nm.

**Supplementary Figure 3:**
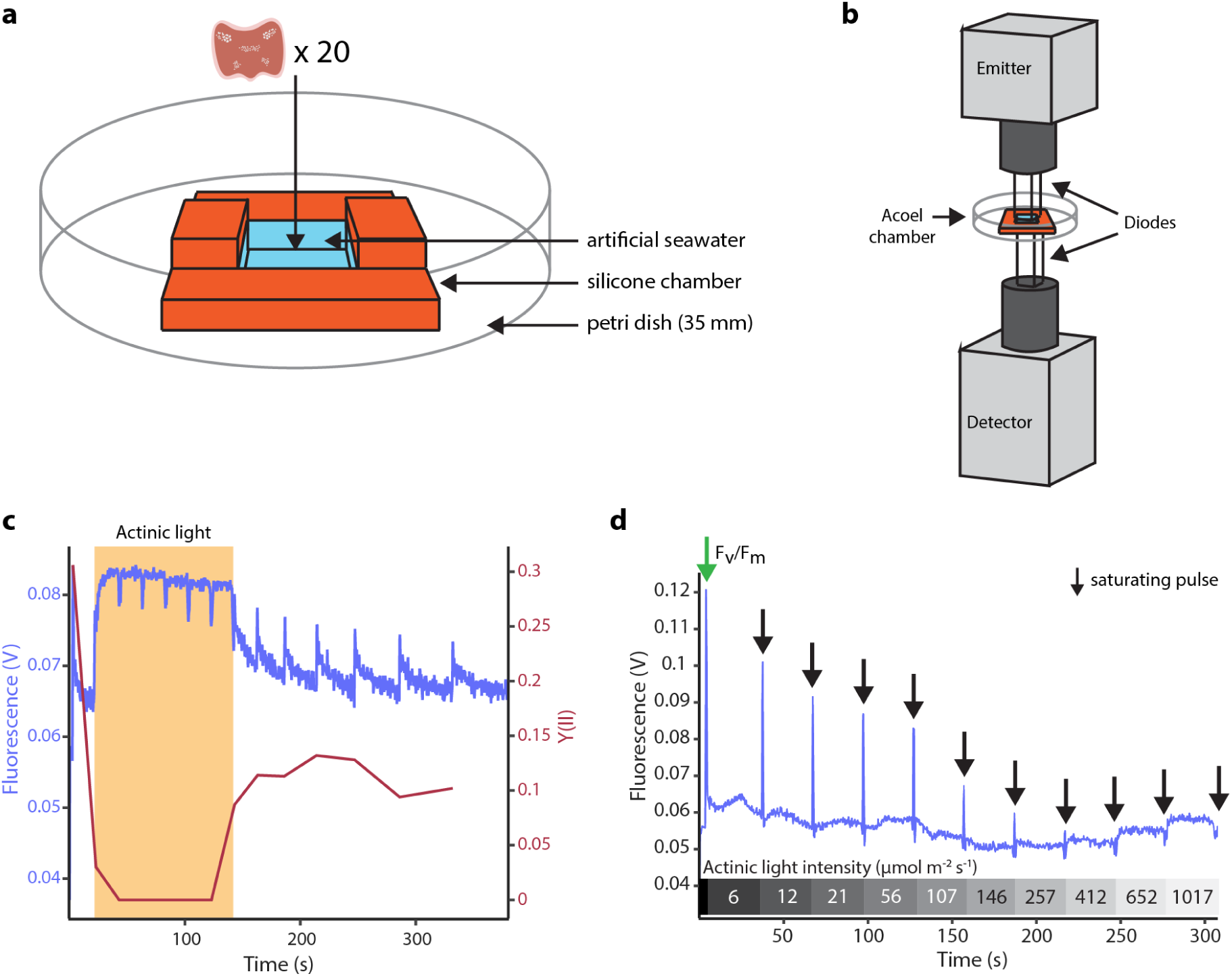
Measuring photosynthetic efficiency of symbiotic algae within the acoel host using PAM fluorometry. **a** Custom chamber designed for constraining acoels during PAM measurements. The chamber, made from a silicone mold, has an inner space of 5 × 6 × 2mm. Two taller walls (4 mm in height) prevent the emitter diode from touching the water. The chamber is mounted on a 35 mm petri dish. For each experiment, 20 tails are transferred to the chamber right before measurements, and the chamber is filled by ASW. **b** The vertical setup of the PAM system is used to mount the acoel chamber between the emitter (top, covering the chamber’s inner area) and the detector (bottom, in direct contact with the chamber). **c** Fluorescence measurement showing DCMU treatment (20 μM) inhibits algal photosynthesis. After the first saturating pulse to measure F_v_/F_m_, a constant actinic light is turned on (orange background) causing the fluorescence to rapidly increase to the maximum level and the Y(II) to drop to zero, showcasing PSII blockage **d** Raw fluorescence trace of tails at 0 hpa. The bar at the bottom shows the actinic light intensity during the fluorescence measurements. The arrows point at saturating pulses, and the size of the fluorescence peak is used to calculate Y(II) at different actinic light intensities. Beyond 146 μmol photons m^-2^ s^-1^ the Y(II) could not be calculated due to the fluorescence signal during saturating pulses being within the background steady-state fluorescence.

**Supplementary Figure 4:**
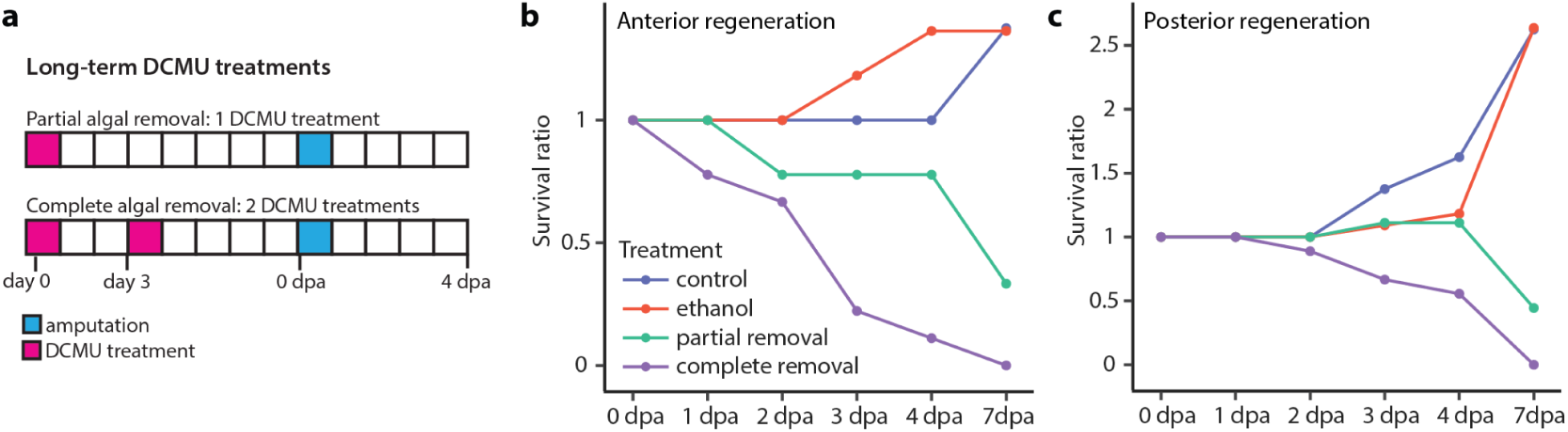
Long term DCMU treatments lead to acoel death. **a** Schematic showing the long-term DCMU treatment time course, with each square representing a day. The day of acoel amputation is marked in blue, while the days of DCMU treatments are marked in magenta. Each DCMU treatment lasts for 24 hr. A single DCMU treatment leads to partial algal removal, whereas two treatments are sufficient to remove algal cells completely. As DCMU is resuspended in ethanol, we treat animals with matching amounts of ethanol as controls. **b,c** Survival ratio of animals from no-treatment control, ethanol-treated control, partial algal removal, and complete algal removal groups during anterior (**b**) and posterior (**c**) regeneration. In control groups, animals undergo fission, which results in more animals at the end of the experiment..

**Supplementary Figure 5:**
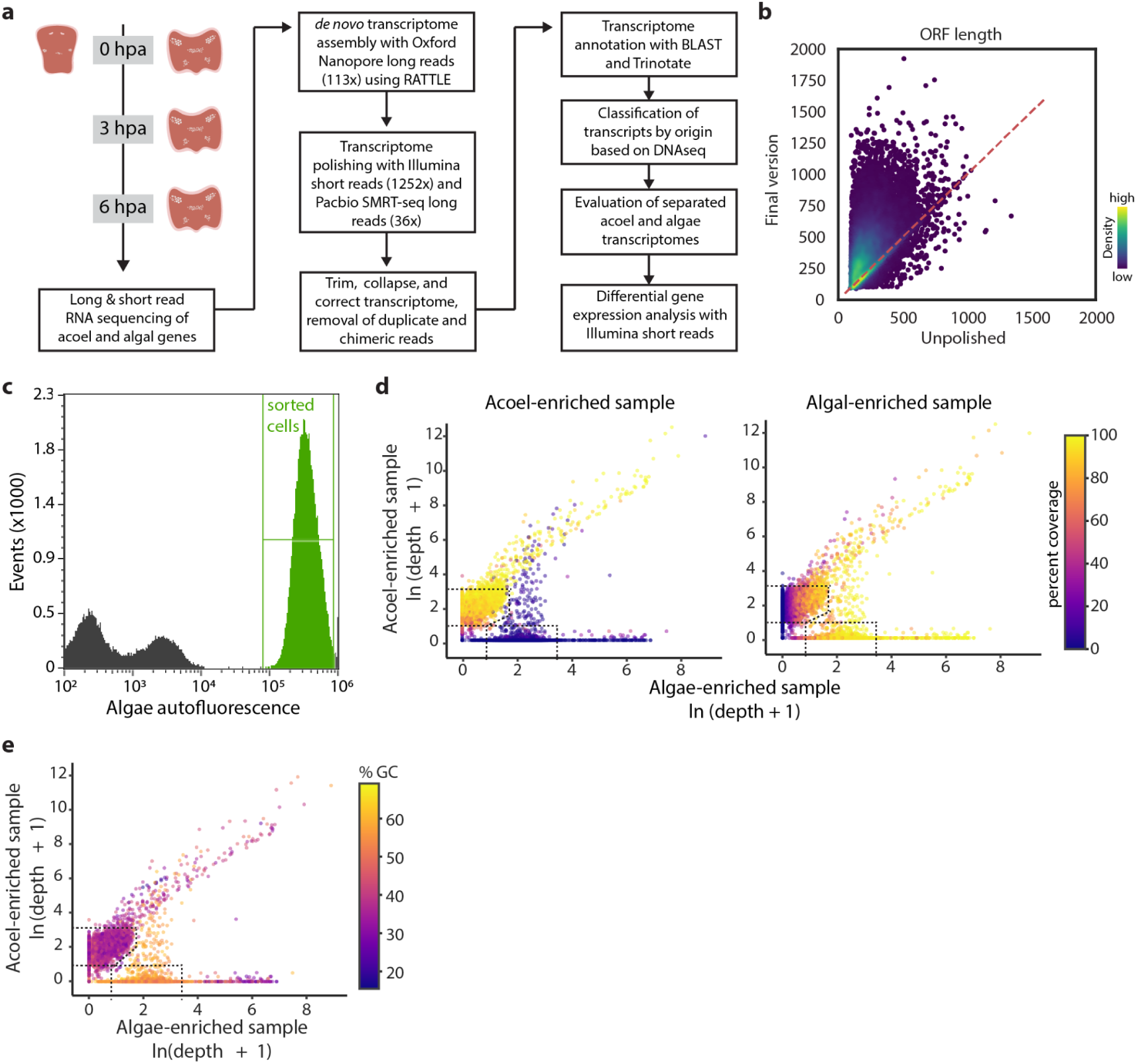
*De novo* transcriptome assembly and annotation. **a** The workflow to assemble the acoel and algal transcriptomes *de novo*. **b** Density plot comparing ORF lengths before and after polishing. 77.8% of transcripts have longer ORF lengths after polishing. **c** Histogram of algal autofluorescence used for gating algae during flow sort (green gate). **d** Percent coverage, the fraction of transcripts sampled by DNA reads of a particular sample, overlaid on the sequencing depth plot for the acoel-enriched (left) and alga-enriched samples (right). **e** Percent GC content of each transcript overlaid on the sequencing depth plot. In general, algal transcripts have high GC contents compared to acoel transcripts.

**Supplementary Figure 6:**
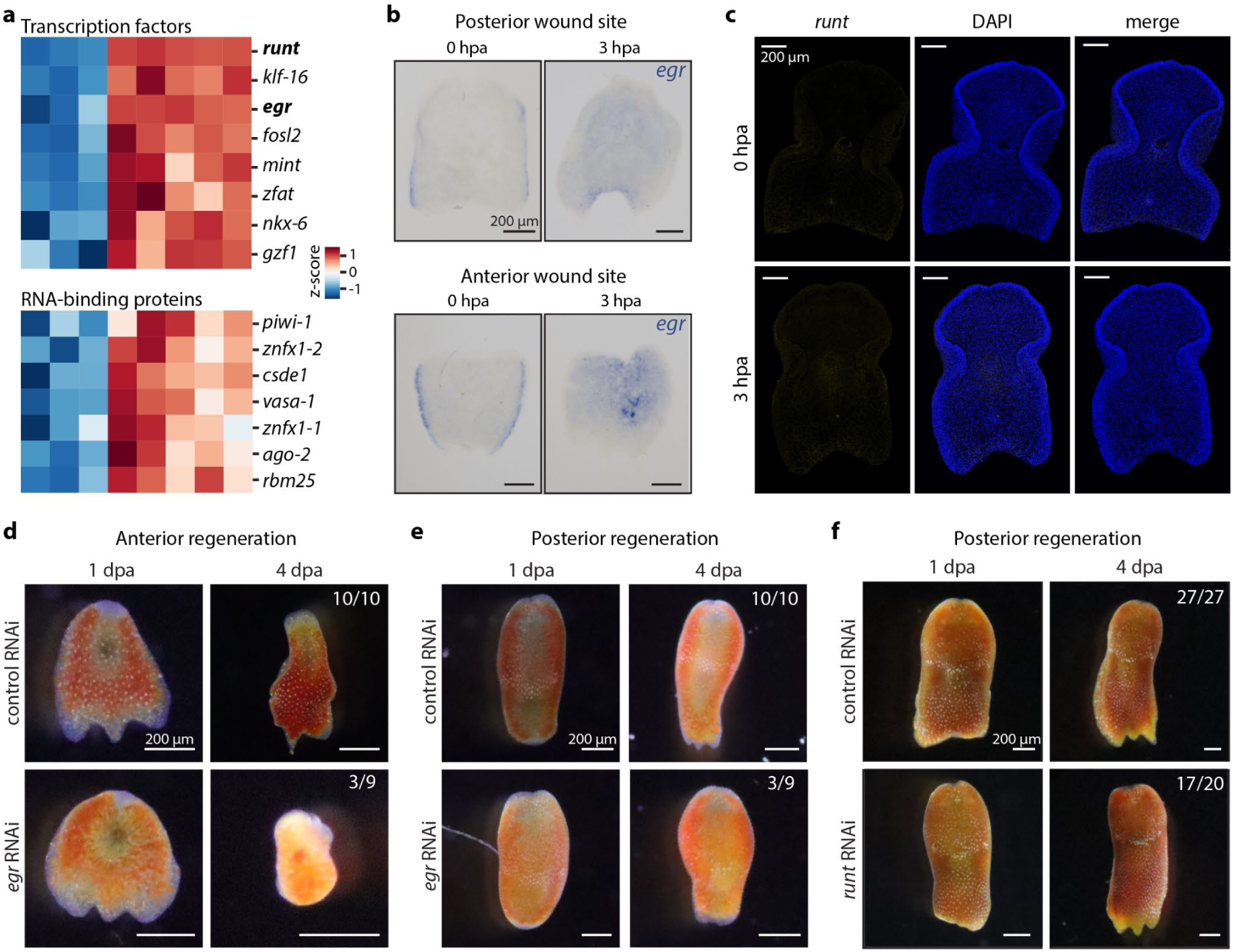
Acoel wound responses. **a** Heatmap of selected acoel regulatory genes upregulated after injury. Top: transcription factors; bottom: RNA-binding proteins. **b** *egr* expression detected with whole-mount in situ hybridization at both anterior and posterior wound sites at 3 hpa. **c** *runt* expression is not detected by HCR at 0 and 3 hpa at the posterior wounds. **d, e** Approximately one-third of *egr* RNAi tail (**d**) and head (**e**) fragments fail to regenerate. Some tail fragments also experienced tissue degradation making the phenotype difficult to interpret. Numbers represent the number of acoels that regenerated similarly to the images shown out of the total number of animals examined. **f** Most *runt* RNAi tail fragments regenerate three-lobed tails by 4 dpa. Numbers represent animals that regenerated out of the total evaluated animals.

**Supplementary Figure 7:**
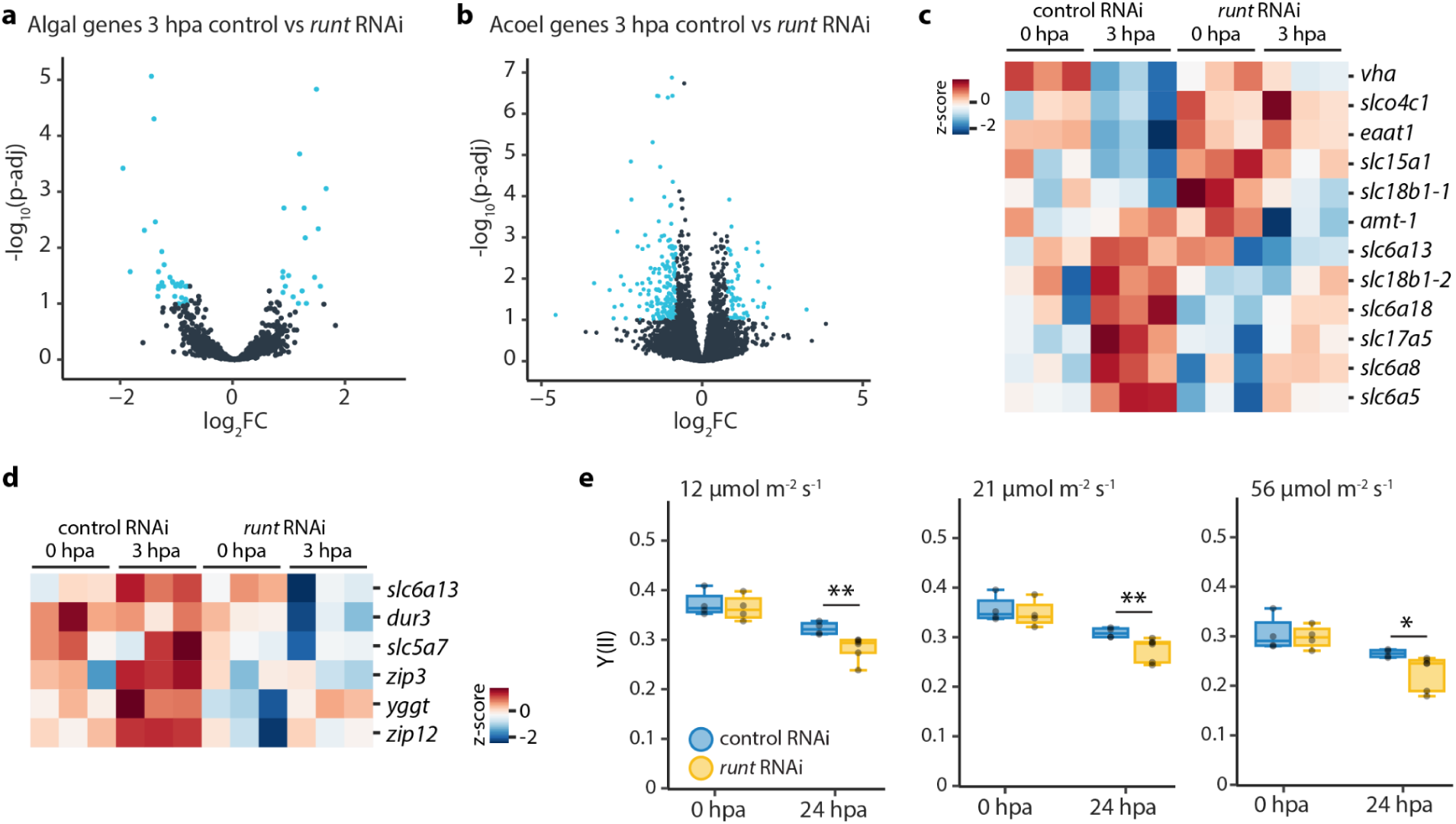
Effects of *runt* RNAi on regeneration responses. **a,b** Volcano plots of algal (**a**) and acoel (**b**) genes at 3 hpa control vs *runt* RNAi comparison. Differentially expressed genes are shown in blue (log_2_FoldChange ≧ 0.8 or ≦ -0.8, p-adj ≦ 0.1). **c,d** Heatmaps of acoel (**c**) and algal (**d**) transporters affected by *runt*. **e** Y(II) of *runt* RNAi and control tails at 0 and 24 hpa with different intensities of actinic background light. Both *runt* RNAi and controls show a decrease in Y(II) at 24 hpa compared to their counterparts at 0 hpa. Y(II) of *runt* RNAi at 24 hpa compared to control RNAi is significantly lower (**pval ≦ 0.05, *p-val ≦ 0.1, t-test). Each treatment consisted of four replicates measured once, and each replicate contained twenty tails.

## Supplementary Tables

**Supplementary Table 1.** Summary of RNA sequencing runs for transcriptome assembly.

|  | Nanopore | Pacbio | Illumina |
| --- | --- | --- | --- |
| Number of raw reads | 4,205,343 | 681,514 | 285,726,698 |
| Number of filtered reads | 3,224,900 | 679,170 | 280,739,473 |
| Average read length (bp) | 1,129 | 1,701 | 143 |
| Transcriptome coverage | 113x | 36x | 1,252x |

**Supplementary Table 2. Gene annotations for acoel and alga transcriptomes.** Species origin and gene names used in this study are provided in separate columns.

**Supplementary Table 3.** Comparison of transcriptome quality. *H. miamia* transcriptome is obtained from Gehrke, et al. ^29^, *Isodiametra pulchra* and *Tetraselmis suecica* obtained from NCBI database. The transcriptomes presented in this study have longer transcripts, fewer chimeric transcripts, and a higher fraction of transcripts with predicted ORFs.

|  | *C. longifissura*<br>(this study) | *Tetraselmis sp.*<br>(this study) | *H. miamia* | *Isodiametra pulchra* | *Tetraselmis suecica* |
| --- | --- | --- | --- | --- | --- |
| % complete + fragment BUSCOs | 79.2 | 54.9 | 97.2 | 94.9 | 39.2 |
| % transcripts without ORF | 10.3 | 4.46 | 12.6 | 72.6 | 17.4 |
| auN ORF length (aa) | 519 | 491 | 753 | 882 | 554 |
| auN transcript length (bp) | 1,889 | 1,863 | 2,384 | 2,503 | 1,805 |
| mean % GC | 40.2 | 57.6 | 36.1 | 45.9 | 57.8 |
| mean ORF length (aa) | 359 | 366 | 345 | 121 | 319 |
| mean transcript length (bp) | 1,514 | 1,547 | 1,294 | 831 | 1,266 |
| N50 ORF length (aa) | 486 | 443 | 521 | 602 | 449 |
| N50 transcript length (bp) | 1,781 | 1,719 | 1,727 | 1,561 | 1,495 |
| Number of transcripts | 13,313 | 7,270 | 20,951 | 66,257 | 11,645 |
| ORF auN/mean ratio | 1.45 | 1.34 | 2.18 | 7.29 | 1.74 |
| Total size (bp) | 20,149,048 | 11,251,596 | 27,117,555 | 55,082,050 | 14,749,373 |
| transcript auN/mean ratio | 1.25 | 1.2 | 1.84 | 3.01 | 1.43 |

**Supplementary Table 4. Oligo sequences used in this study.** Oligo pools for HCR to detect *runt* expression, primers for cloning gene fragments used in RNAi (*runt*, *egr*) and ISH (*egr*, *pc2*) experiments, and custom oligos for RNA-seq experiments are listed with their sequences.

## Supplementary Data

**Supplementary Data 1: Fasta file for the assembled acoel transcriptome.**

**Supplementary Data 2: Fasta file for the assembled algal transcriptome.**

